# THE RIBOREGULATION MECHANISM OF HUMAN SERINE HYDROXYMETHYLTRANSFERASE IS ROOTED IN AN ALLOSTERIC SWITCH

**DOI:** 10.1101/2023.03.17.532973

**Authors:** Sharon Spizzichino, Federica Di Fonzo, Chiara Marabelli, Angela Tramonti, Antonio Chaves-Sanjuan, Alessia Parroni, Giovanna Boumis, Francesca Romana Liberati, Alessio Paone, Linda Celeste Montemiglio, Matteo Ardini, Arjen J. Jakobi, Alok Bharadwaj, Paolo Swuec, Gian Gaetano Tartaglia, Alessandro Paiardini, Roberto Contestabile, Serena Rinaldo, Martino Bolognesi, Giorgio Giardina, Francesca Cutruzzolà

## Abstract

RNA can directly control protein activity in a process called riboregulation; only a few mechanisms of riboregulation have been described in detail, none of these being characterized on structural grounds. Here we present a comprehensive structural, functional, and phylogenetic analysis of riboregulation of cytosolic serine hydroxymethyltransferase (SHMT1), the enzyme interconverting serine and glycine in one-carbon metabolism. We show that the RNA modulator competes with polyglutamylated folates and acts as an allosteric switch, selectively altering the enzyme’s reactivity vs. serine. In addition, we identify the tetrameric assembly and a flap structural motif as key structural elements necessary for binding of RNA to eukaryotic SHMT1. The results presented here suggest that riboregulation may have played a role in the evolution of eukaryotic SHMT1 and the compartmentalization of one-carbon metabolism. The findings also provide insights for RNA-based therapeutic strategies targeting this cancer-linked metabolic pathway.

## INTRODUCTION

RNA binding proteins represent a complex group of proteins, often interacting with RNA to form stable or transient ribonucleoprotein complexes through canonical RNA binding domains (RBDs)^1–4^. Over the past few years, it became clear that proteins may also bind RNA with non-canonical RBDs and that such unconventional RNA binding proteins are often involved in responding to environmental and physiological cues^2^. Intriguingly, unconventional RNA-binding properties have been reported for enzymes involved in many aspects of cellular metabolism, from glycolysis to citric acid cycle, gluconeogenesis, amino acid biosynthesis, fatty acid oxidation, lipid metabolism, and nucleotide metabolism^5,6^. The RNA:enzyme interaction is often exploited to modulate the expression of target RNAs, as in the case of aconitase/IRP1 regulating ferritin mRNA, or GAPDH controlling IFNγ mRNA translation^7,8^. The overall perspective on the biological significance of RNA:protein interactions has been lately shifting from a protein-centric approach, where proteins regulate the RNA, to a more complex view, whereby RNA can directly affect protein structure and function, in a process called riboregulation.

Through riboregulation, RNA can control protein aggregation^9^, intracellular localization^10^, interaction with ligands and enzymatic activity^11^. In the few mechanistic studies available on metabolic enzymes, binding of RNA and substrates were shown to be mutually exclusive events, as observed for thymidylate synthase (TYMS) and dihydrofolate reductase (DHFR), which cannot bind their cognate mRNAs in the presence of substrates^12,13^. The same concept applies to human glycolytic enolase I (ENO1), where several RNA molecules compete with the substrate molecule for binding to the enzyme active pocket^14^.

Riboregulation was shown to control the activity of human enzyme serine hydroxymethyltransferase (SHMT), a pyridoxal 5’-phosphate (PLP) dependent enzyme involved in the folate-dependent serine/glycine interconversion in one-carbon (1C) metabolism, a key network responsible of nucleobases biosynthesis and redox balance (Figure 1 A-B) ^15^. In humans, the two main SHMT isoforms, encoded by the two genes *SHMT1* and *SHMT2,* are localized in the cytosol and mitochondria, respectively^16^. Interaction of the cytosolic SHMT1 with the 5’UTR of the *SHMT2* mRNA transcript (UTR2, 206 nt long) lowers the expression of the mitochondrial isoform SHMT2 (Figure 1B)^15,17,18^. Remarkably, the same interaction riboregulates SHMT1 enzymatic function in a selective manner by inhibiting the serine to glycine cleavage reaction, but not the reverse reaction, *i.e.* serine synthesis (Figure 1B) ^15^. Therefore, RNA not only binds to but also selectively riboregulates SHMT1. By dynamically exploiting these two biological effects of the RNA:SHMT1 interaction, cells can finely tune the serine-glycine metabolism across different cellular compartments^19^.

**Figure 1.**
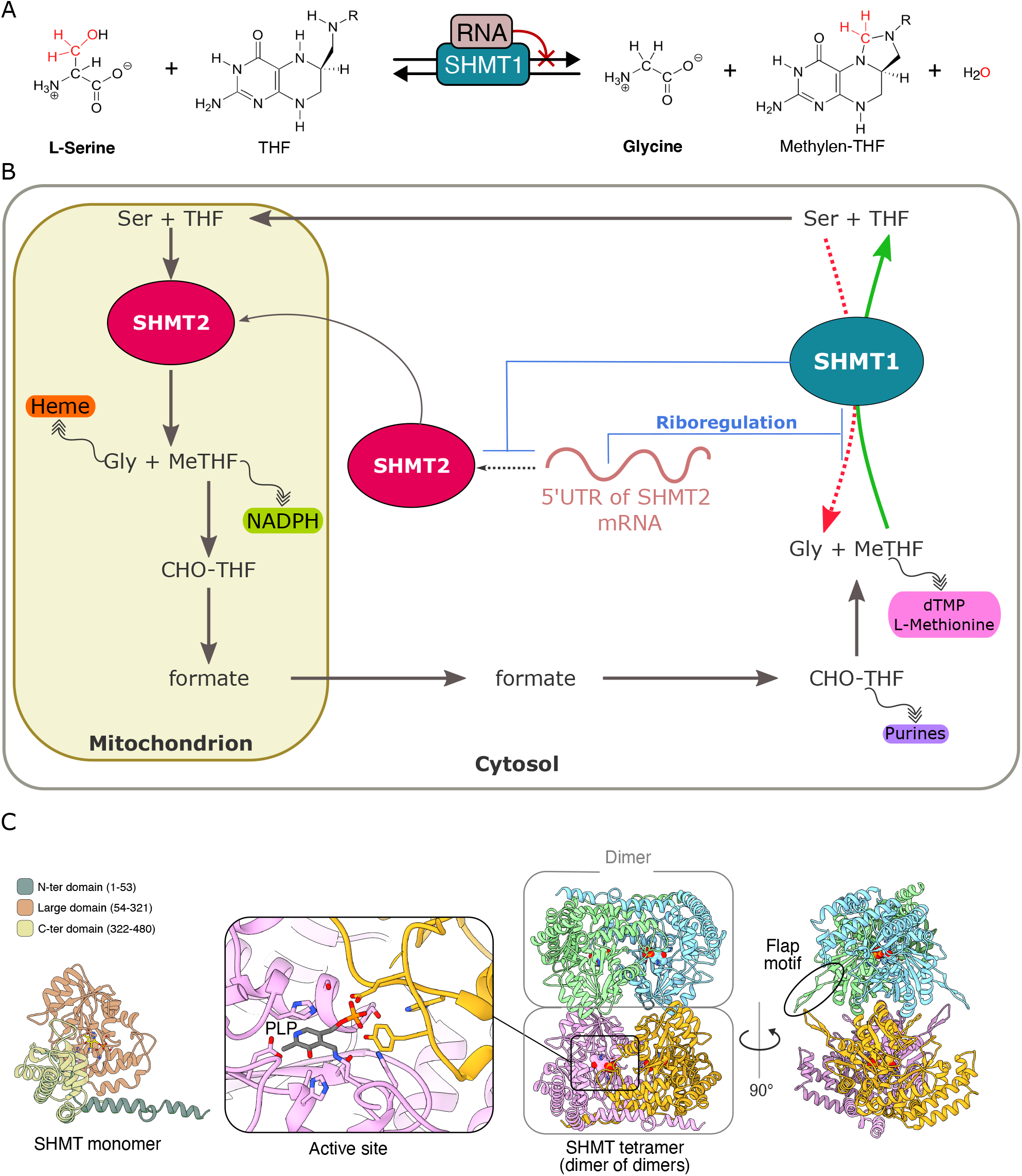
Structure, catalytic activity and riboregulation of SHMT. **A)** Scheme of the reversible reaction catalysed by SHMT1. RNA riboregulates the enzyme by inhibiting only the serine to glycine conversion and not the reverse reaction. **B)** The scheme shows the reactions catalysed by SHMT1 and SHMT2 in the context of the folate metabolism cycle taking place between the cytosol and mitochondria^58^. As shown in Panel A, The 5’UTR of SHMT2 mRNA riboregulates SHMT1 enzymatic activity; in turn, SHMT1 controls the translation of SHMT2 mRNA ^15^. **C)** Quaternary structure and domain organization of human SHMT. Human SHMT1 is shown with each monomer differently colored (left panel) (1BJ4^32^). The PLP coenzyme (shown in grey) is located in the dimeric interface and therefore the dimer is the minimal catalytic unit. Only one active site (out of the four ones) is highlighted in the figure. The position of the flap motif is highlighted in black.

SHMTs are often highly expressed in cancer cells and their expression correlates with poor prognosis ^20–24^. Despite their clinical relevance, effective small molecule inhibitors of SHMT, able to interfere with cancer cells proliferation and metastasis formation *in vivo,* are still missing^18,25^. This couples with the poor structural characterization of the complex regulatory mechanisms, including riboregulation, controlling SHMT activity.

Here we present the cryo electron microscopy (cryo-EM) structures of human SHMT1 in its free- and RNA-bound states. The structural data are complemented by *in vitro* RNA-binding and inhibition studies based on selected RNA constructs and substrates, and by evolutionary analyses. Our results unravel the SHMT1 RNA-binding mode and highlight a conformational transition in two distinct subunits of the tetrameric enzyme. Taken together, our structure-based mechanistic study shows that a RNA hairpin is the primary element binding SHMT1 and that riboregulation promotes a selective allosteric inhibition of the enzyme catalysed serine-to-glycine reaction.

## RESULTS

### Cryo-EM structure of SHMT1: an open conformation of the active site is populated in the absence of substrates

SHMT catalyses the reversible conversion of serine to glycine, through a coupled reaction transforming tetrahydrofolate (THF) into N5, N10-methylene-THF (CH2-THF) (Figure 1A).

The catalytically active unite pf SHMT is a dimer, containing two active sites located at the dimer interface and each containing one PLP molecule (Figure 1C)^26–28^. Eukaryotic enzymes assemble into a tetramer (dimer of dimers); solution studies revealed that each subunit within the quaternary assembly is in equilibrium between two conformations: open and closed^16,29–31^. The open conformation displays a more flexible and solvent exposed active site, which becomes more rigid and buried in the closed state. The open/closed equilibrium was proposed to be differently affected by binding of the serine or glycine substrates ^29^. Remarkably, the only crystal structure available of human SHMT1 shows a perfectly symmetric tetramer, with each subunit being in the closed conformation (PDB ID 1BJ4^32^).

The structure of the unliganded SHMT1 was determined here by single particle cryo-EM at 3.2 Å resolution (Figures 2, S1 and supplementary movie 1); no symmetry was imposed during the data processing. The four subunits are indicated as A, B, C and D (chain ID of the PBD 8C6X coordinates), with the A-B subunits forming one dimer and the C-D subunits forming the other (arranged in the tetramer as A-B|C-D). Surprisingly, in the SHMT1 tetrameric structure here reported, only chains A and D nicely match the closed conformation of the previously published crystal structure and display a defined density for the bound PLP and of the residues interacting with the cofactor (all the Cα of chain A and D of the cryo-EM structure superpose with the crystal structure (1BJ4) with RMSD values calculated on 462/465 of Cα atoms of 0.89 Å and 0.83 Å, respectively). On the contrary, chains B and C lack electron density within the cofactor binding region and their C-terminal domains are shifted (to a maximum distance of 4.1 Å; RMSD values calculated on 462/465 Cα atoms are 1.11 Å for chain B and 1.14 Å for chain C)^32^ (Figure 2). Both observations suggest that the active sites in B and C subunits are in an open conformation; therefore, we will refer to these open states as B* and C* (Figure 2). The active site open conformation was never clearly observed within the crystallographic structures ^27,32,33^ possibly due to lattice restraints or to conformational selection during crystal growth. The cryo-EM results show that the active sites of SHMT1 can assume, in the absence of substrates, both the closed and open conformations, the latter previously inferred only from solution studies^29,30,34^. The SHMT1 tetrameric assembly, showing the location of the open and closed subunits (A-B*|C*-D state) is shown in Figure 2.

**Figure 2.**
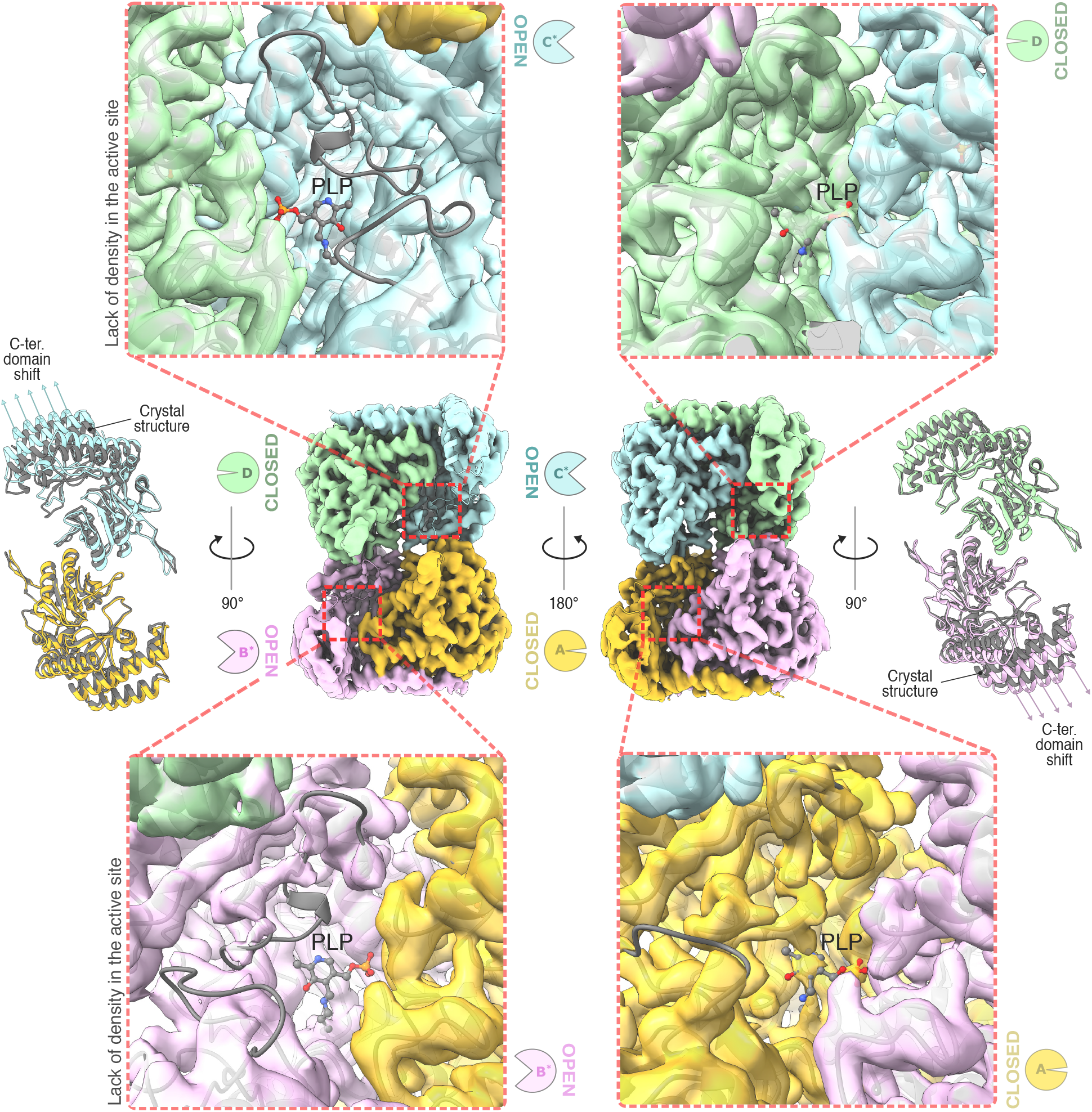
Cryo-EM structure of unbound SHMT1. Sharpened electron density map of the unbound SHMT1 (PDB:8C6X). Chain B and C are in an open conformation, whereas chain A and D are in a closed conformation. **Upper and lower panels**: focus on the active site of each subunit. The subunits with an open conformation (on the left) show a lack of electron density in the active site region. PLP is shown in grey in ball and sticks**. Far left and right panels**: structural superposition of each subunit of the unbound SHMT1 (different colours) with the individual subunits seen in the crystal structure (in grey: PDB 1BJ4^32^). Only the residues belonging to the large domain were used in the alignment to highlight the shift of the C-terminus domain of the subunits in the open conformation.

### RNA binding induces a conformational reorganization in the conformation of SHMT1 subunits

The structure of the SHMT1:RNA complex was determined by cryo-EM at 3.5 Å resolution, in the presence of an RNA fragment corresponding to the first 50 nucleotides of UTR2 (hereinafter UTR2_1-50_). Also in this case, no symmetry was applied during the data processing. The structure of the complex displays one RNA molecule bound to the SHMT1 tetramer, thus achieving 1 RNA molecule per tetramer stoichiometry (Figure 3A).

**Figure 3.**
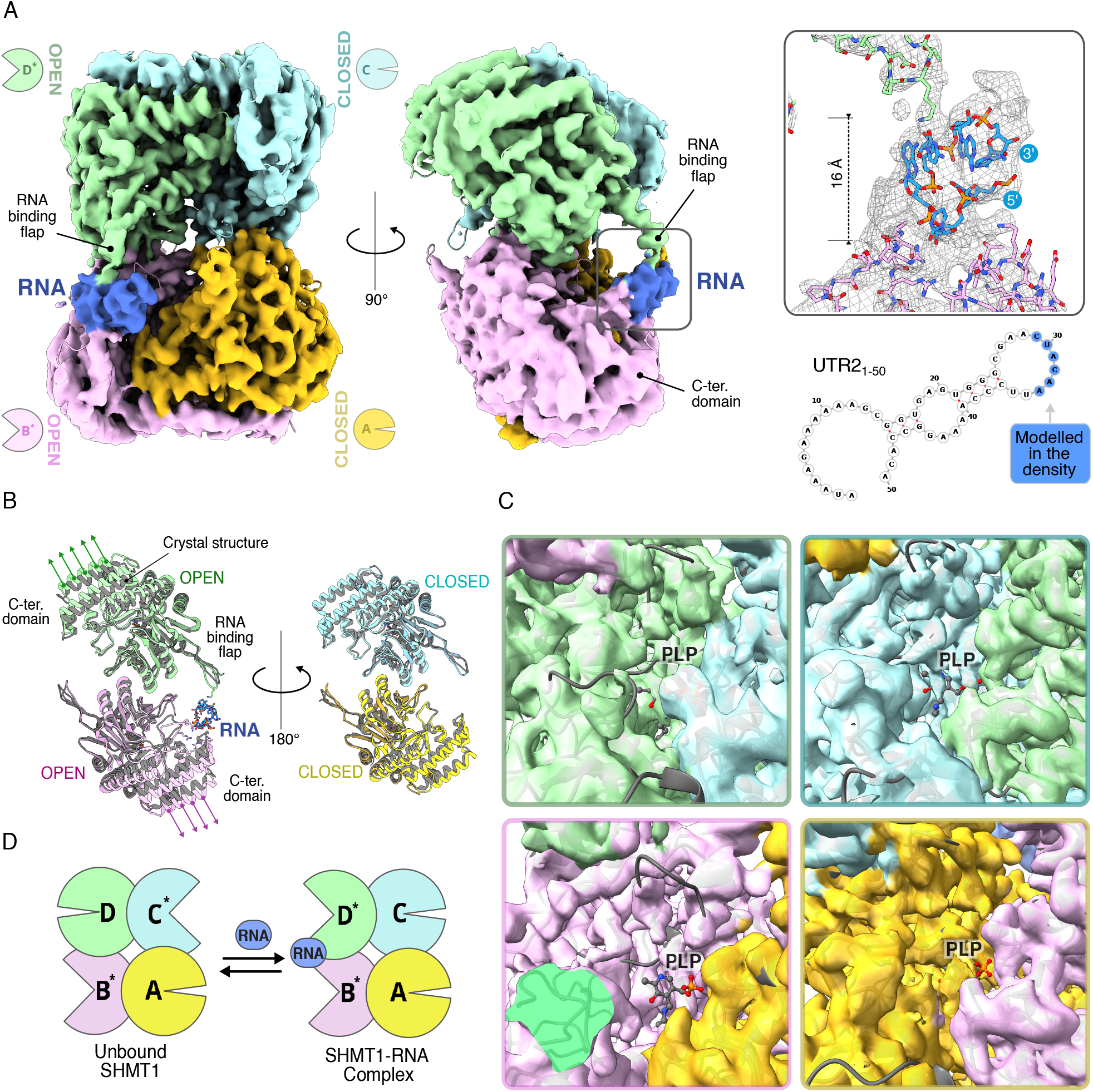
Structural features of the SHMT1-RNA complex. **A)** Sharpened electron density map of the complex (PDB 8A11). The four subunits and the RNA (blue) are rendered in different colours. The final model of the 6 nt built in the RNA extra density and the predicted secondary structure (Vienna RNAfold web server: <u>RNAfold web server</u>) for UTR2_1-50_ are shown on the right. **B)** Structural superposition of each subunit of the SHMT1-RNA complex (different colours) with the individual subunits seen in the crystal structure (in grey: PDB 1BJ4^32^). **C)** Electron density in the active site region for the four subunits. The PLP is labelled, an extended region of density centred around His-148, that makes an aromatic stacking interaction with the coenzyme, is not visible or poorly defined in both the subunits involved in RNA binding (B* and D*). **D)** Scheme of the conformational topology of the unbound SHMT1 and the RNA complex. The wide aperture corresponds to the open conformation.

The RNA fragment lies in front of one active site (subunit B), contacting the C-terminal domain of subunit B and one flap motif of the adjacent subunit D belonging to the opposite dimer (Figure 3A and supplementary movie 2). RNA binding thus relies on the SHMT1 tetrameric assembly and on the presence of the flap motif, a β-hairpin structure composed of 13 amino acids (273-VKSVDPKTGKEIL-285), whose function in eukaryotic SHMT was never fully clarified, although a recent study tried to define its role^35^. The SHMT1:RNA cryoEM structure finally assigns a clear purpose to this motif, which will be referred to as the “RNA binding flap”.

With respect to the crystallographic structure, the two SHMT1 subunits (B and D) contacting the RNA display a shift of their C-term domains (RMSD values calculated superimposing the cryo-EM SHMT1:RNA structure and the crystal structure (1BJ4) on 462/465 of Cα atoms are 1.08, 1.22, 1.02 and 1.11 Å for chains A,B,C and D respectively) (Figure 3B), which correlates with a lack of electron density in their active site clefts, showing that the open and closed conformations of the subunits are present also in the SHMT1:RNA complex (Figures 3B, 3C and S2). In both cryo-EM structures, the two open subunits belong to distinct dimers; the topology of the open and closed subunits observed in the RNA-bound tetrameric SHMT1 (A-B*| C-D*) is however different from that of the unbound form, as shown in Figure 3D. In RNA-free SHMT1, the active sites in the open conformation appear to be on average more dynamic than those in the complex, according to B-factor analysis (Figure S3A), suggesting that RNA binding may also affect the overall flexibility of the protein. Interestingly, RNA binds to the protein regions that oscillate the most; normal mode analysis shows that the C-term and flap regions are the most mobile, as the distance of the Cα-backbone of the RNA binding flap and the C-terminal domain is constantly fluctuates between 20 and 27 Å (Figure S3B and supplementary movie 3).

The cryo-EM structures of SHMT1 (unbound and bound to RNA) show that the enzyme can populate distinct quaternary states, each showing a different combination of the active sites in the open/closed conformation (Figure 3D); this observation suggests an RNA-dependent allosteric control of the enzyme. The presence of allostery was previously observed in solution and in structural studies of SHMT in the presence of substrates. In solution titration calorimetric studies, the ligand 5-formylTHF (an unreactive THF derivative) was found to bind with a stoichiometry of 2/tetramer, and to display negative cooperativity^31^. In addition, in the crystal structure of mouse SHMT in complex with 5-formylTHF and glycine (1EJI^26^), the folate is not bound to all four available active sites, suggesting that, also in this case, binding of the folate to one dimer reduces the affinity for the other dimer (negative cooperativity)^26^.

Based on the above observations, we thus hypothesize that binding of RNA may decrease the conformational mobility of SHMT1 in solution and select the quaternary state of SHMT1 tetramer (A-B*| C-D*) observed here.

### SHMT1 binding to the RNA is non-sequence specific and requires a hairpin loop

The specificity of RNA-protein interactions generally relies on the flexibility of the ribose-phosphate backbone, on the RNA secondary and tertiary structure, and on the interactions formed by purine and pyrimidine bases. In the SHMT1:RNA complex, the observed RNA-density was only partly defined, likely due to the high flexibility of the nucleic acid. This extra density accounted for six nucleotides belonging to the predicted hairpin region of the UTR2_1-50_ (C29UACAA34), which were modelled and refined in real space with good geometry and are at H-bonding distance from Lys279 (Figure 3A). Such binding pose is strongly dependent on the size of the RNA hairpin, and, on one hand, it is not achievable by single stranded RNA, too small to fill the gap between the C-term domain and the flap. On the other hand, a double stranded RNA would be too large to fit the density.

This structure-based hypothesis is supported by binding experiments in solution, showing that SHMT1 binds with higher affinity to single stranded RNA oligos containing a hairpin loop (Figure S4 A and B). Double stranded RNA does not bind SHMT1 and lower binding affinities were found for RNA molecules with loops smaller or larger than 14 nt (Figure S4A). In addition, UTR2_1-50_ and its reverse complement RNA, with different nucleotide sequences but sharing a similar secondary structure, bind with similar affinities (Figure S4C and D), suggesting that, in the SHMT1:RNA complex, the interaction does not require a specific sequence, as often observed in other RNA binding proteins, including TYMS and DHFR^6,36–38^.

### Mapping the RNA binding site reveals an RNA binding hub on SHMT1 tetramer

RNA binding to SHMT1 is achieved by a combination of spatial constraints and electrostatic interactions. Four lysine and three arginine residues contact or are near to the RNA docking site (Lys157, Lys158, Lys279, Lys282, Arg372, Arg393 and Arg397), suggesting that electrostatic interactions play a leading role in guiding RNA through the binding process (Figure 4A). On the C-term domain, RNA docks to a positively charged cleft (Figure 4A), exposing arginine residues Arg372, Arg393 and Arg397. In keeping with such structural arrangement, mutation of key positively charged amino acids lowers or abolishes RNA binding (Figure 4B). Notably, mutation of residues Lys157, Lys158 ^15^ and Lys282 (this work) that are not in direct contact with the structured RNA hairpin mapped in the density, reduce the binding affinity similarly to the mutation of interacting residues (e.g. the Arg393-Arg397 double mutant and the Lys279 single mutant) (Figure 4B; Figure S5A and B). These observations suggest that the part of the RNA oligo (50 nt long) extending from the hairpin can interact with the protein region surrounding the main RNA binding site, possibly adapting several different conformations. It is interesting to note that close inspection of our cryoEM raw data highlighted the presence of lower-resolution 3D classes with blurred extra densities extending from the RNA-binding site (Figure S1A). This indicates a relatively high degree of flexibility of the RNA oligo, whose main docking site is located between SHMT1 C-term domain and the flap, thus achieving the most stable (and thus selected *in silico)* SHMT1:RNA assembly (Figure S1A).

**Figure 4.**
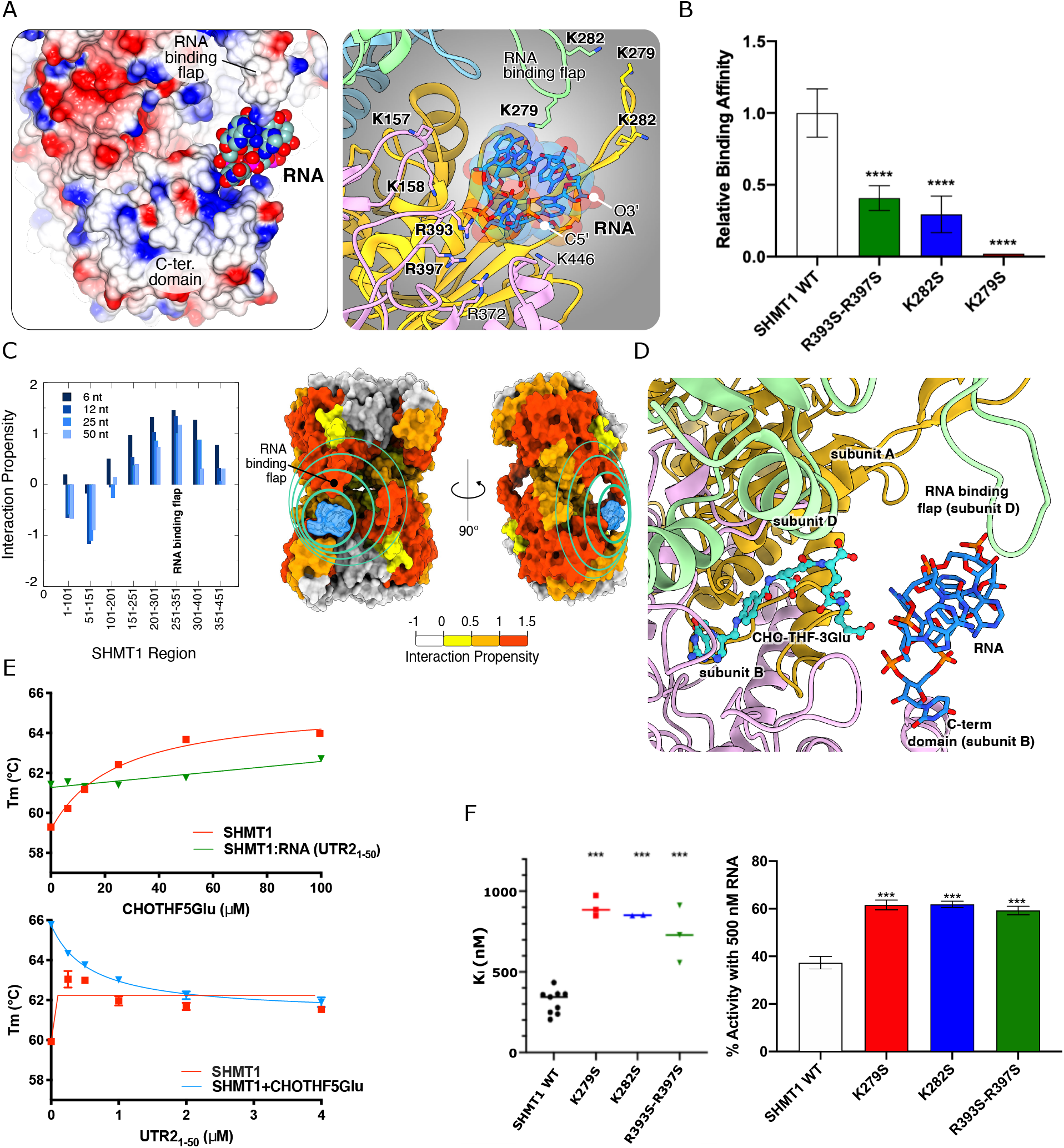
Mapping the RNA-binding region. **A)** Left panel: Electrostatic surface potential of the RNA binding region. RNA inserts into a positively charged cleft and is fastened against the RNA binding flap. Right panel: Positively charged residues that are in contact or in proximity to the RNA. **B)** Binding affinity of the mutants of key SHMT1 residues interacting with RNA (UTR2_1-50_), all the experiments were performed three times (n=3). The binding affinity of SHMT1 WT and mutants (left graph) was obtained by EMSA (shown in Figure S5). A value of 1 was assigned to the WTSHMT1-RNA interaction, which showed the best affinity, and all the other mutants (K279S, K282S and the double mutant R393S-R395S) were compared to it. The mutation of the key residues involved in the RNA interaction affects the inhibitory effect exerted on the protein. Statistical analysis is performed on three independent experiments. **** P ≤ 0.0001. **C)** The interaction propensity (correlated with binding affinity) of RNA for different SHMT1 regions. Interaction simulations of UTR2_1-50_ fragments of different length were run against 100-residue long SHMT1 regions with a 50-residue overlap. Mapping the results with different colours on the surface of SHMT1 shows a large region of interaction, centred around the RNA binding hub, and extending around the tetrameric interface like a belt. **D)** The RNA and THF binding sites on SHMT1 are located in the same region. The crystal structure of rabbit SHMT1 in complex with 5-formyl-THF-3Glu (CHO-THF-3Glu) (PDB: 1LS3) was aligned with the SHMT1-RNA complex (different colours, PDB: 8A11). The RNA is coloured in light blue, and CHO-THF-3Glu in cyan. **E)** Effect of RNA on the binding of CHO-THF-5glu probed by DSF experiments. The changes in the Tm of the protein-folate or protein-RNA complexes are plotted as a function of ligand concentration. Data were fitted with an equation describing a hyperbolic saturation behaviour. **F)** The inhibitory effect of UTR2_1-50_ on the mutants R279S, R282S and the double mutant R393S-R395S, compared to the WT protein. Left panel: All the experimental data, acquired in at least three independent experiments, were calculated using equation 2 (see Materials and Methods) and fitted by using a sigmoidal function. The plotted Ki are a mean of the Ki obtained in each individual experiments and are 315±25, 903±37, 852±2 and 733±102 nM for SHMT1 WT, K279S, K282S and R393S-R397S respectively. Right Panel: the plot shows the residual activity of the protein(s) when using 500 nM RNA. The obtained activity are 37.3±2.6, 61.6±2.0, 61.9±1.3 and 59.3±1.7 for SHMT1 WT, K279S, K282S and R393S-R397S respectively. Statistical analysis is performed on three independent experiments. *** P ≤ 0.001.

The above structural and mutagenesis data suggest that the RNA hairpin docks to one region, whereas the single-stranded part of RNA flexibly extends along the SHMT1 surface to the adjacent potential RNA binding site(s) on the same side of the tetramer (Figure 4A). We hypothesize that the RNA molecule samples the protein surface through an ensemble of binding modes centred around the observed 6-nucleotide density, in a region that we may define as “RNA-binding hub”; thus, the RNA binding free energy landscape may be characterized by multiple minima of comparable energy.

To confirm this hypothesis, we performed *cat*RAPID^39^ calculations of RNA-protein interaction propensity using different stretches of the RNA_1-50_ oligo and running the calculations against protein regions of 100 residues with a 50-residue overlap. Interestingly the results show a broad region of interaction centred around the SHMT1:RNA interaction site (in particular the flap) and extending in both directions at the level of the tetrameric interface (Figure 4C).

### RNA competes with the binding of folates

Structural superposition of the RNA binding site in human SHMT1 and the folate binding site from rabbit cytosolic SHMT in complex with 5-formyl-THF-3Glu (hereinafter CHO-THF-3Glu)^40^, confirms that the RNA and folate binding sites fall in the same region of the enzyme (Figure 4D).

The binding mode of RNA to human SHMT1 here reported resembles that observed for the negatively charged polyglutamate chain of the folate substrate. In the rabbit SHMT:CHO-THF-3Glu complex, the folate species adopts multiple conformations in each of the two occupied binding sites, the polyGlu tail being electrostatically bound to surface cationic residues^40^. Polyglutamylated folates represent the physiological form that traps folates within the cell^41,42^ enhancing their interaction with cognate enzymes by order of magnitudes (from 5.9 μM to 20 nM)^31^.

Our structural data on the SHMT1:RNA complex indicate that both RNA and folate binding require a tetrameric enzyme, since they bind to two subunits (B and D in Figure 4) belonging to two different dimers. Interestingly, only the polyGlu tail stretches out of the folate binding site and interacts with the flap residues 273-285.

To further explore the similarities between RNA and polyGlu folates binding, we analyzed by Differential Scanning Fluorimetry (DSF) the effect of RNA on the association of a pentaglutamylated form of folate (CHO-THF-5Glu) to SHMT1. As shown in Figure 4E by titrating SHMT1 with CHO-THF-5Glu, an increase in Tm is observed (ΔTm=+4.68 ± 0.12°C), suggesting binding of folate species to the protein under these conditions. On the contrary, no significant Tm change is observed when the binding experiment is run starting from the SHMT1:RNA complex. Accordingly, addition of RNA to the SHMT1:CHO-THF-5Glu complex, leads to a decrease in Tm (ΔTm: −3.89°± 0.35 °C), suggesting that the RNA oligo is able to displace CHO-THF-5Glu.

### Riboregulation of SHMT1: mechanism and selectivity

Our structural and solution binding results suggest that the SHMT1:RNA interaction requires a tetrameric assembly, the flap motif and identify an RNA binding hub located nearby the folate binding site. To address the question of whether the molecular determinants of RNA binding and of riboregulation map to the same region of the protein, we firstly established that mutation of the positively charged residues Lys279, Lys282, Arg393 and Arg397 identified in the RNA binding hub also significantly decreased the ability of RNA to inhibit SHMT1 (Figure 4F), while the intrinsic catalytic activity of mutated SHMT1 was not significantly affected (Figure S5C). Next, we demonstrated that the tetrameric state is important for riboregulation by showing that the serine cleavage activity in a bacterial SHMT, which is dimeric, is unaffected by the presence of RNA (Figure S6A).

To better analyze the effect of RNA on the enzymatic activity, we first confirmed that the UTR2_1-50_ RNA fragment inhibits the SHMT1 catalyzed serine cleavage reaction (see Figure 1A), but has no effect on the reverse glycine-to-serine conversion (Figure 5A). By increasing the concentration of RNA in the Ser→Gly reaction, we observed an increase in the apparent Km for THF (Figure 5B). This finding and our data presented above, confirms that a partial competitive or hyperbolic mechanism is operative, as previously suggested^15^. This type of inhibition is often characteristic of allosteric inhibitors^43–45^. Moreover, substrate inhibition, typically observed at high THF concentration and attributed to the formation of a resting enzymatic state (the enzyme– glycine–THF dead-end complex)^46^, becomes less evident after pre-incubation with RNA (Figure 5B), suggesting again a significant influence of the nucleic acid on THF binding.

**Figure 5.**
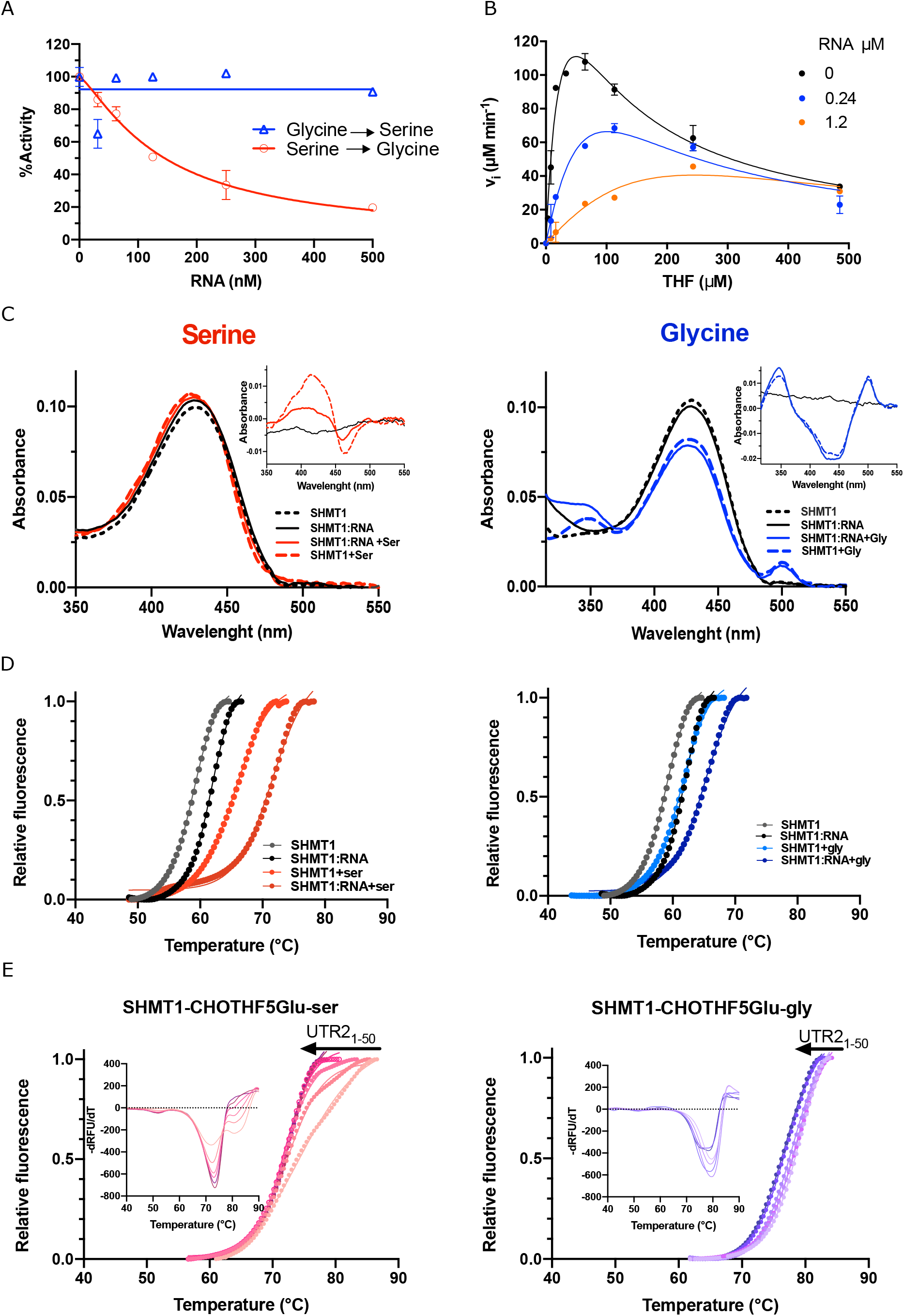
Effect of amino acid substrates on the activity of SHMT. **A)** Effect of UTR2_1-50_ on human SHMT1. The serine cleavage activity of SHMT1 in the presence of UTR2_1-50_ was measured. The inhibition constant (Ki) of the UTR2_1-50_ RNA towards human SHMT1 is 141.5 ± 13.7 nM. The inhibitory effect of the RNA was also tested in the reverse reaction (Gly→Ser) catalysed by human SHMT1, confirming that the inhibition seen is selective for the serine cleavage activity of the enzyme. **B)** Inhibition of the SHMT1-catalyzed Serine cleavage Reaction by UTR2_1-50_ RNA. The serine cleavage reaction catalysed by SHMT1 (0.28 μM) was assayed in the presence of 10 mM L-Serine and increasing concentration of THF (3-485 μM) in 20 mM K-Phosphate buffer pH 7,2 at 30°C. The experiment was carried out in the absence of UTR2_1-50_ RNA (Black circles) and by incubating the enzyme in the presence of UTR2_1-50_ RNA 0.24 e 1.2 μM (blue and orange circles). **C)** Binding of amino acid substrates (Serine and Glycine) is differentially affected by UTR2_1-50_. The spectra of the protein:amino acid complexes were recorded in the presence of 10 mM L-Serine (left panel) and 10 mM Glycine (right panel) without or with pre-incubation of SHMT1 (18 μM) with UTR2_1-50_ (4.5 μM) in 50mM HEPES; 150mM NaCl; 5% glycerol; pH 8 at 25°C. (inset) Differential spectra of SHMT1:amino acids complexes calculated by subtracting the spectrum of the protein alone, or that of the SHMT1:RNA complex. The difference spectra of the enzyme in the presence of RNA alone is also shown for comparison. **D)** DSF experiments probing the effect of serine and glycine (10 mM) on the stability of the SHMT1 and SHMT1:UTR2_1-50_ complex. 0.5 μM of UTR2_1-50_ were mixed with 2 μM SHMT1. **E)** The effect of increasing the amounts of UTR2_1-50_ (0.25-4 μM) on the stability of the ternary complex formed by mixing SHMT1 2 μM:CHO-THF-5Glu 50 μM:Seine or Glycine 10mM) with the indicated amino acid as measured by DSF experiments. The direction of the observed changes upon addition of the RNA is indicated by an arrow. In the DSF experiments, to better show the observed different transitions, the error bars are not shown in the plot.

To explain the differential effect of RNA on the direct (Ser→Gly) and reverse (Gly→Ser) reactions, we then investigated the effect of RNA on the protein environment (around the amino acid substrates) within the active site, using UV-VIS spectroscopy. In the first step of the reaction, the substrate amino acids react with the PLP cofactor to form spectroscopically distinct derivatives (see scheme in Figure S6B). Pre-incubation of SHMT1 with the UTR2_1-50_ RNA, before addition of the amino acid substrate (serine or glycine), significantly alters the formation of serine derivative(s), while leaving those observed after addition of glycine unaffected (Figure 5C and S6A). This shows that, relative to glycine, serine is more sensitive to the RNA-induced changes of the active site structural and chemical environment, and suggests that the RNA-bound enzyme is in a state that selectively hampers productive serine binding within the pocket. Notably, correct positioning of serine Cβ atom in the active site was previously proposed to be required for triggering the transition from the open to the closed conformation in the enzyme subunits^47^.

To gain further insight into the differences between the serine and glycine complexes of SHMT1, we compared their thermal stability through DSF experiments and then assessed the effect of RNA. The binding of serine and glycine has a different stabilizing effect on SHMT1 (ΔTm of +7.0 ±0.1 and +2.7 ±0.1°C, respectively, at 10 mM amino acid concentration) (Figure S7 and Table S1), in agreement with literature data indicating that serine should trigger a conformational change when binding to the open subunits, while glycine should not. A different behaviour of the two amino acids is also observed when they are added to the SHMT1:CHO-THF-5Glu binary complex: in the experiments with serine, an additional transition from a species with Tm 75.7±0.2 °C to a more stable one with a higher Tm (83.8±0.5 °C) is observed at high serine concentration, whereas in the case of glycine an apparent two-state unfolding process is observed, regardless of glycine concentration (Tm 78.6±0.1 °C) (Figures 5D and S8). In both cases, extremely stable ternary complexes (SHMT1:CHO-THF-5Glu:amino acid) are formed: the observed increase in stability is larger than the sum of the effects of individual ligands (Figure S8 and Table S1), suggesting that a change in the allosteric state of the enzyme is likely taking place when both substrates (amino acid and folate) are present. Notably, when the poly-Glu tail is not associated to the folate (as for THF, CHO-THF, Lometrexol) (Figure S9), the hyper stabilization effect is lost for both serine and glycine, strengthening the hypothesis that this effect is linked to a change of the enzyme quaternary allosteric state requiring the extended glutamate tail that can contact the flap motif.

The binding of UTR1-50 RNA to SHMT1 increases the Tm of about 3°C, while resulting in a destabilization of the SHMT1:CHO-THF-5Glu binary complex (Figures 4E and 5D and Table S1). We then characterized the influence of RNA on the stability of the binary (SHMT1:amino acid) and ternary (SHMT1:CHO-THF-5Glu:amino acid) complexes. A differential effect of RNA is already present in the binary complexes (Figure 5D, Table S1), but the most striking result is observed in the experiments on the ternary complexes with the two substrates (Figure 5E, Table S1). In the case of serine, the second transition mentioned above disappears and the denaturation profile reverts to an apparent two-state transition; on the contrary, the denaturation profile of the ternary complex with glycine is not significantly affected. The ternary complex with glycine is thus resistant to the effect of RNA. These results clearly show that RNA is able to modify the allosteric states populated by serine, while having a marginal effect on those populated by glycine.

Altogether, we observed RNA-dependent changes both in the active site and in the allosteric state transition when measuring the interaction with serine, but not with glycine, providing the molecular bases of the selective inhibitory effect on SHMT activity.

### The SHMT1 tetrameric assembly and the flap co-evolved with the compartmentalization of one-carbon metabolism and riboregulation

Our previous data^15,19^ suggested that an evolutionary pressure was exerted on eukaryotic SHMT by the need to compartmentalize and differentially regulate one-carbon metabolism in cytosol and mitochondria. Such pressure might have resulted in the evolution of SHMT quaternary structure (i.e., dimer-to-tetramer assembly and flap acquisition) that, in turn, allowed the onset of SHMT1 riboregulation. Since it is not clear when exactly these structural fingerprints of eukaryotic SHMTs riboregulation emerged during Evolution, we carried out a systematic phylogenetic analysis starting from the Archaean eon, to assess whether the tetrameric quaternary structure and the flap motif underwent a coevolutionary process (Figures 6, S10, S11 and Table S2).

**Figure 6.**
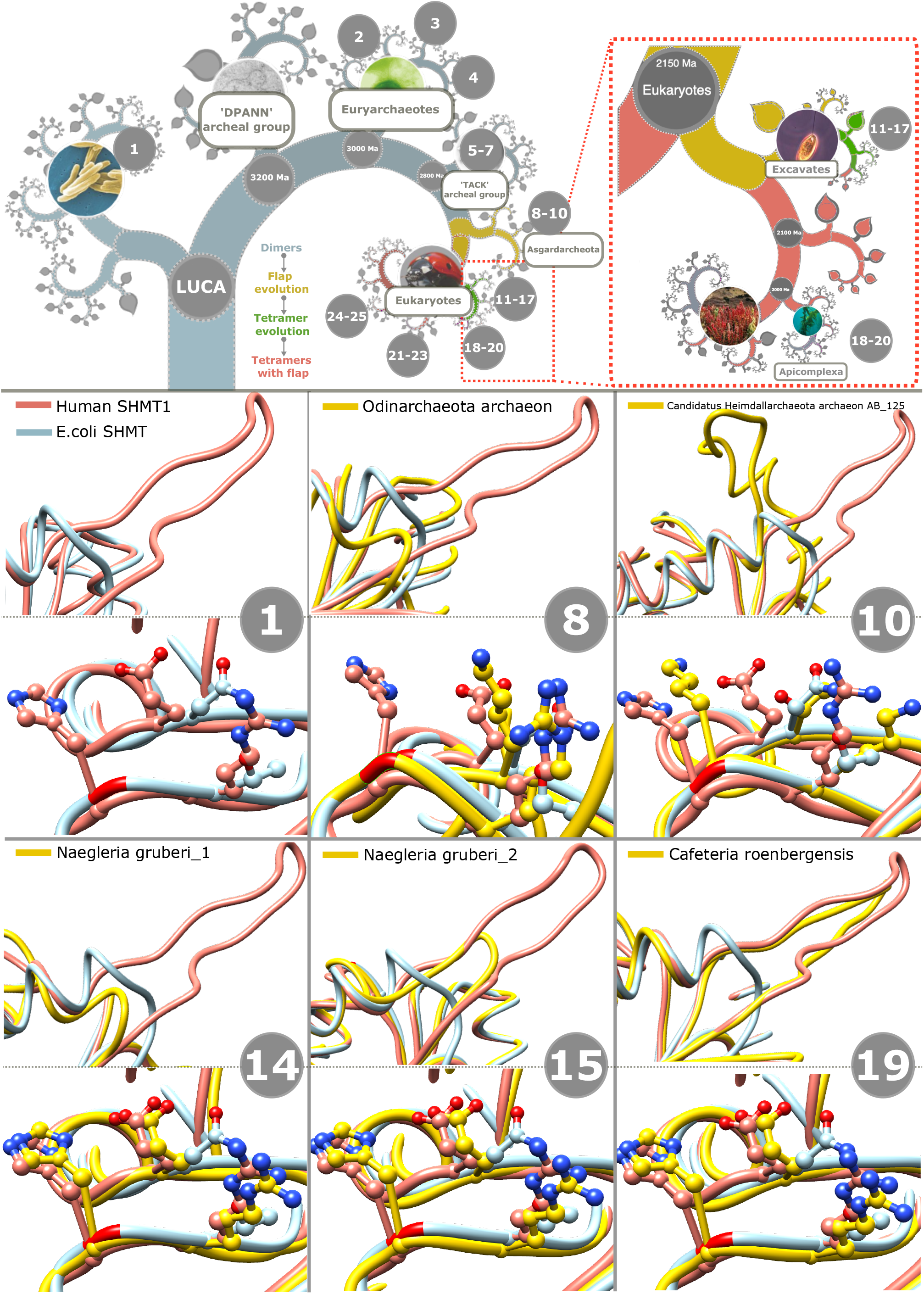
Evolutionary analysis. **Upper panel:** Phylogenetic tree and evolution of SHMT quaternary features. The Flap motif (yellow) has started to evolve in the superphylum of Asgardarchaeota, before that divergent point the protein was dimeric (light blue). The tetramer (green) has started to evolve in the Unikonta subdomain. The tree has been generated by OneZoom.org/life^59^. **Lower panel:** structural superimposition of human SHMT1 (pdb 1bj4, pink), SHMT from E.coli (1DFO, light blue) and the one from the analysed organism (yellow). The presence or absence of the flap motif and the tetramerization residues (His 135, Arg 137, and Glu 168 in human) is highlight Further details are reported in Table S2.

Starting from the *Euryarchaeota,* where all the analyzed SHMTs lack the flap motif and the three main tetramer-stabilizing residues (His135, Arg137, and Glu168)^18,32^, and moving along the evolutionary tree to the superphylum of *Asgardarchaeota,* we found that dimeric SHMTs feature a flap motif, analogous to that of the eukaryotic SHMTs, only in the *Heimdallarchaeota* subphylum. The observation of the flap motif in dimeric SHMTs from *Asgardarchaeota* indicates that the flap appeared during evolution before the tetrameric assembly, since it is conceivable that it was already present in the common ancestor of *Eukaryota* and *Asgardarchaeota* (Table S2). The reason for the evolutionary emergence of this feature is unclear, but it might be related to an archetypically acquired RNA-binding function, which became progressively essential along with the complex re-compartmentalization of the metabolic fluxes between cytoplasm and the nascent mitochondria. Hence, the tetrameric quaternary structure could have subsequently arisen between 2.7 and 2.1 billion years ago, in the evolutionary branch connecting *Eukaryota* and *Asgardarchaeota,* to strengthen the possible regulatory role exerted by RNA.

When analysing SHMTs from the *Eukaryota,* we observed that in the *Unikonta* subdomain all SHMTs analysed have features typical of a tetrameric assembly and host flap motives, while in the *Bikonta* lineages we found a more complex scenario: some SHMT isoforms are “bacterial-like” structures, i.e., dimeric without the flap-motif (e.g., *N. fowleri, P. cosmopolites, P. Vivax),* whereas those tetramer-competent (e.g., *E. gymnastica, S. oncopelti, A. castellani, D. purpureum)* show either structured flap motifs, or no-flap-motif at all.

Interestingly, this reversal to a “bacterial-like” SHMT structure in some *Bikonta* lineages is always related to adaptive lifestyles (i.e., anaerobic symbionts and parasites) affecting the mitochondria. Many members of the *Bikonta* (e.g., amitochondriates) lack mitochondria, while others retain a mitochondrial organelle in a greatly modified form (e.g., a hydrogenosome or a mitosome)^48^, or with peculiar evolutionary and functional characteristics (e.g., Plasmodium^49–51^).

Thus, the phylogenesis of SHMT suggests that SHMT1 tetrameric assembly and the flap motif, required for RNA:protein interaction, co-evolved with the compartmentalization of one-carbon metabolism between cytoplasm and mitochondria, linking riboregulation to a sophisticated control of metabolic routes.

## DISCUSSION

Metabolic reprogramming allows cells to adapt to intrinsic or extrinsic cues through high flexibility in nutrient acquisition and utilization^23,52^: riboregulation of metabolic enzymes shows that the cellular response may also be controlled by RNA molecules.

The riboregulatory potential of modulatory RNAs may dynamically change based on their structure and conformation, availability of substrates and cofactors, and intracellular localization of the partners. It appears clear that the SHMT1:RNA complex is characterized by a high degree of conformational flexibility of both the associated partners, in keeping with the concept that RNA structural dynamics plays a key role in guiding its cellular functions ^53,54^. It also confirms that, in the case of SHMT1, the RNA inhibitory effect exploits the dynamics of the target enzyme, which populates different states in solution.

Our structural and functional results demonstrate that RNA competes with the polyglutamylated form of the substrate folate, and selects an allosteric state of the enzyme which has significant consequences on the mechanism of the Ser→Gly conversion reaction. Indeed, both the interaction with serine at the level of the active site, and the effect of this amino acid on the ensemble of states sampled by the protein in solution, are significantly altered in the presence of RNA. These effects are not observed with glycine, providing a plausible mechanism for the selectivity of riboregulation on SHMT serine cleavage activity (see Scheme in Figure 7).

**Figure 7.**
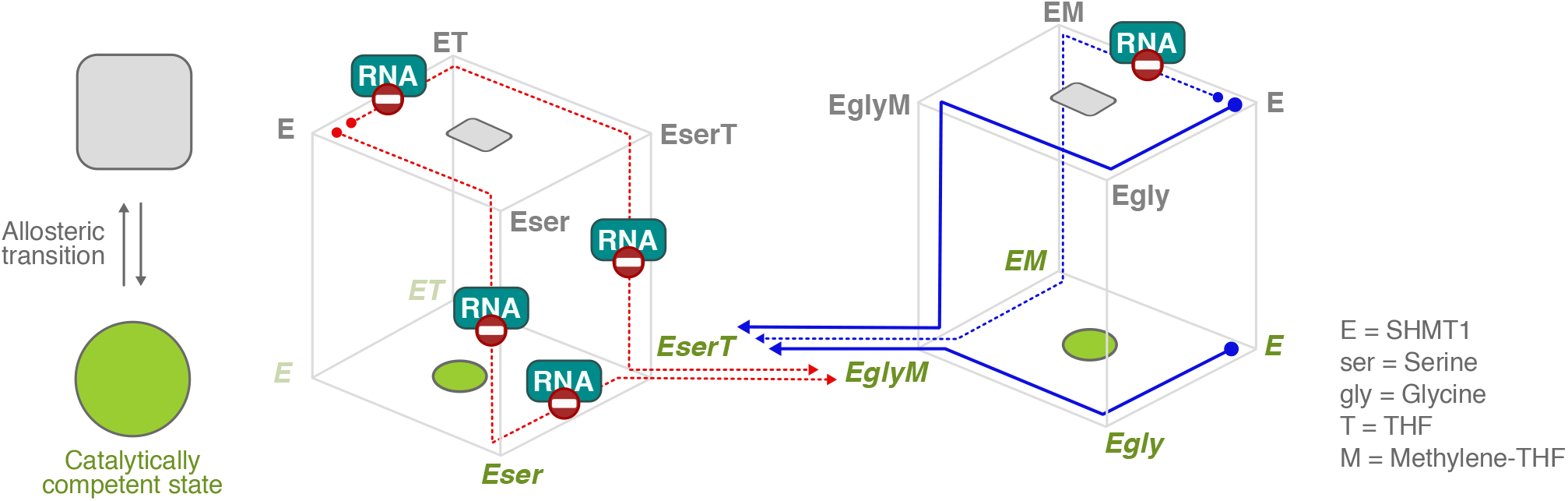
Simplified scheme of the proposed riboregulation mechanism of SHMT1. The scheme summarizes the possible catalytic routes for Ser→Gly (on the left in red) and Gly→Ser (on the right in blue) conversion. All species possibly populated in the reactions are shown, with species known to be less populated represented in light colour^30,31,34^. The enzyme (E) binds randomly to the two substrate^58^ serine or glycine (ser and gly) and the folate substrate THF or methylene-THF (T and M, respectively). The enzyme can exist and bind substrates in at least two different allosteric states (gray square and green circle, upper and lower routes). Binding of ser or formation of the ternary complexes (EserT and EglyM) induces an allosteric transition to a catalytically competent state. In the scheme, the reaction routes shown to be affected by RNA are indicated as dashed lines (red and blue for serine and glycine, respectively). As depicted in the scheme, while RNA affects several steps of the reaction with serine, it has a more limited influence on the reaction with glycine, thus providing a plausible mechanism for the selective inhibition of the serine cleavage reaction of SHMT1.

Our results also indicate the possible effects of riboregulation on the compartmentalization of one-carbon metabolism. In cancer cells, we have shown that the consequence of increasing the level of RNA species that bind to SHMT1 (i.e. the SHMT2 mRNA) is the separation of glycine and serine utilization between the cytosolic and mitochondrial compartments, through the two enzyme isoforms ^19^. In mitochondria, SHMT2 catalyses the Ser→Gly reaction, and sustains the production of one carbon units to ultimately yield formate that is then exported to the cytosol; for this reason, the folate mitochondrial pool should be kept high to meet the metabolic demand of THF by the SHMT2 serine cleavage reaction. Based on the results presented here, we propose that riboregulation of SHMT1 has two parallel and physiologically relevant effects, (i) it favors the glycine to serine conversion in the cytosol to increase the serine levels available for mitochondrial 1-C metabolism, and (ii) it can limit THF entrapment in the dead-end complex of the cytosolic enzyme (as seen in the inhibition studies presented here), thus simultaneously increasing the THF levels available for use in the mitochondria by the same pathway.

In agreement with our observations on the mechanistic effects of RNA binding, the phylogenetic analysis indicates that the RNA-binding structural motifs (tetrameric assembly and the flap motif) have evolved in parallel with the eukaryotic cell and its need to regulate and compartmentalize metabolic functions. Thus, RNA may act as a dynamic metabolic “switch” not only by controlling gene expression, but also by directly affecting the activity of selected targets either by competing with substrates and/or by shifting the equilibrium of allosteric states, as shown in this study.

Our results may contribute to clarify a more fundamental and fascinating issue, i.e. how RNA molecules might have concurred in the emergence of new metabolic abilities and to the shaping and regulation of metabolic routes. RNA:metabolic enzyme pairs might have co-evolved to control the fate of selected metabolites and the direction of metabolic fluxes, as seen in the case of SHMT, thus suggesting a possible scenario explaining how the complexity of metabolism might have raised during evolution, an issue that is a matter of intense debate in the scientific community^55–57^. Finally, exploiting the molecular details of SHMT1 riboregulation also paves the way to the design of regulatory RNAs to be employed as drugs for therapeutic intervention in cancer and other pathological states, targeting SHMT1 and possibly other metabolic enzymes known to host RNA-binding functions.

## MATERIALS AND METHODS

### Materials availability

Chemicals and reagents Serine, Glycine, bovine serum albumin (BSA), trombine, diethyl pyrocarbonate (DEPC), and all the salts were obtained from Sigma-Aldrich. Tetrahydrofolate and (6S)-5-formyl tetrahydrofolate (monoglutamylated forms) were kindly provided by Merck & Cie (Schaffhausen, Switzerland). All other chemicals were used in the enzymatic assays purchased from Sigma-Aldrich.

UTR2 _1-50_ used for the cryo-EM experiments was purchased from Eurofins Genomics (Germany GmbH), instead the UTR2 _1-50_ and UTR2 _1-50_ Rv used for *in vitro* experiments was purchased from biomers.net GmbH 2023. UTR2 loop length 8nt and UTR2 loop length 28nt were purchased from Sigma-Aldrich.

### Protein expression and purification

Wild-type and mutant (K282S, R393S-R397S, K279S) SHMT1 genes were cloned into a pET22b(+) vector (Novagen) and expressed as N-terminal histidine-tagged fused protein in *Escherichia coli* (BL21-DE3). Bacterial cultures were grown at 37 °C in Luria-Bertani (LB) liquid medium supplemented. Purification was performed as previously described by Spizzichino et al^69^. Briefly, bacterial pellets were then resuspended in lysis buffer (20mM Hepes pH 7.2; 150 mM NaCl; 5% glycerol) and lysed by ultrasonic disruption on ice. Soluble proteins were purified by IMAC on a Ni^2+^-His- Trap column (Cytiva, Chicago, IL, USA). SHMT1 eluted with 300 mM imidazole. Fractions containing pure protein were pooled, and imidazole was removed with desalting columns PD10 (Cytiva, Chicago, IL, USA), the histidine tag was then removed by proteolytic digestion with 1U/mg of thrombin (SIGMA). The digestion mixture was then loaded again on the IMAC on a Ni^2+^-His-Trap column (Cytiva, Chicago, IL, USA) and SHMT1, which now binds with low affinity, was eluted with 100 mM imidazole. The protein was injected into a Superdex 200 column (16/600; GE Healthcare) and eluted as a tetramer with the following buffer, 20mM Hepes pH 7.2; 150 mM NaCl; 5% glycerol. Protein concentration was determined by measuring the absorbance at 280 nm and applying the Beer-Lambert law (SHMT1 ε_280_ 47565 M^-1^cm^-1^). Samples were frozen in liquid nitrogen and stored at −20 °C.

### Cryo-EM grid preparation and data acquisition

Fragmentation of the 206 nt UTR2 was necessary to reduce the RNA molecule mobility and to increase the homogeneity of the single particle on the grids. The first 50 nt of the UTR2 were selected for complex formation since this fragment showed the highest binding affinity to SHMT1 and was still able to decrease enzyme activity *in vitro* (UTR2_1-50_: AUAAAGAAAAAAGCGGUGAGUGGGCGAACUACAAUUCCCAAAAGGCCACA) (Figures S4B and S4D).

Purified SHMT1 protein at 0.3 mg/ml concentration in 150 mM NaCl, 20 mM Hepes pH 7.2 buffer, was incubated with UTR2_1-50_ RNA at a 1:5 protein:RNA molar ratio. A 3 μl drop of the sample was applied to glow-discharged Quantifoil 1.2/1.3 300 mesh Cu grids for 20 s and blotted before plunge-freezing in liquid ethane. Grid vitrification was performed using a Vitrobot Mark IV (Thermo Fisher Scientific) at 100% humidity and 4 °C.

A total of 5,450 movies were collected on a Talos Arctica microscope (Thermo Fisher Scientific) operating at 200 kV on a Falcon 3EC camera in electron counting mode. The dataset was automatically acquired using EPU software (Thermo Fisher Scientific). The pixel size was set to 0.889 Å/pixel. Each movie consists of 40 frames with a total accumulated dose is 40 e^-^/Å^2^. Applied defocus ranged between −0.5 and −2.5 μm.

### Single-particle data processing

Movies were motion corrected and dose weighted with MotionCor2 implemented in Relion 3.1^70^. CTF estimation was performed with CTFFIND-4.1.13^71^ using the sums of power spectra from combined fractions. Images were processed using RELION 3.1^70^. A first subset of 10 representative micrographs was subjected to LoG autopicking. After 2D classification, classes with low background were selected and used as models for a 2D-reference based Autopicking on the whole dataset. A total of 4.9 million initial particles were picked and then subjected to multiple rounds of 2D classification. After generating a low-pass filtered initial model from the SHMT1 crystallographic structure (PDB ID: 6FL5), selected class averages with high resolution features (0.9 million particles) were subjected to a first round of 3D classification with C1 symmetry to remove remaining junk particles. of the remaining 0.7 M particles, a second round of 3D classification with C1 symmetry revealed classes with an extra-density located at one out of the four nucleotide-binding sites of SHMT1. Despite the symmetry of SHMT1 homotetramer, we avoided applying any kind of symmetry bias. Selected classes containing similar extra-densities or the putative RNA-freeform were separately grouped and subjected to further 3D classification to select a subset of homogeneous particles. Final 3D classes selected contained 94,430 and 73,523 particles for the RNA-bound and the RNA-free states respectively. These were subjected to masked 3D auto-refinement, CTF refinement and Bayesian polishing in Relion. The estimated resolution of the final maps, following standard post-processing procedures in RELION^72^, was 3.52 Å for the RNA-bound SHMT1 and 3.29 Å for the RNA-free state. An overall scheme of the image processing pipeline is presented in Figure S1.

### Local scale map sharpening

Local map sharpening was performed using a model-free implementation of the software LocScale (https://gitlab.tudelft.nl/aj-lab/locscale/). The original implementation of LocScale^73^ utilizes information on the amplitude of the radially averaged structure factor computed locally from an atomic model with refined atomic displacement (ADP) factors. In the model-free implementation used here, no prior model information is employed and the radial structure factor is instead approximated by a composite radial profile consisting of a shape-dependent fall-off in the low resolution region (infinity to ~0.1 Å-1) that is defined by the molecular envelope, and a generalised radial structure factor scaled by the local B factor in the Wilson region (~0.1 Å-1 to Nyquist frequency ^74^). The sharpened density map was then constructed by locally applying the scaling procedure in a rolling window of based on this generalised reference as previously described^75^.

### Model building

The initial models of the SHMT1-RNA complex and unbound SHMT1, whereas generated performing a rigid body fit of SHMT1 crystal structure (PDB: 1BJ4 (Renwick, Snell and Baumann, 1998)) by using UCSF Chimera^76^. Each subunit was fitted independently. The initial model was then manually modified and refined in Coot (Emsley et al., 2010) to fit the sharpened map or, in the more dynamic regions, the unsharpened one.

The 6 nucleotide hairpin RNA (CUACAA, belonging to the sequence of UTR2 1-50 AUA AAG AAA AAA GCG GUG AGU GGG CGA ACU ACA AUU CC CAA AAG GCC ACA, was fitted into the extra electron density in Coot as well ^77^. The initial geometry of the RNA hairpin was modelled based on a bacterial RNA structure (pdb: 2JR4 ^78^) and then mutated and refined in the electron density in Coot. The full atomic models of SHMT1-RNA complex and SHMT1 unbound were then subjected 3 cycles of real-space refinement using PHENIX^79^, including global minimization and refinement of atomic displacement parameters, and applying secondary structures, Ramachandran, and nucleic acids restraints ^80^.Full statistics are reported in Table 1.

**Table 1.** Cryo-EM data collection, image processing and model refinement statistics.

| Structure | SHMT1 | SHMT1-RNA |
| --- | --- | --- |
| Microscope | TALOS Arctica |  |
| Voltage (kV) | 200 |  |
| Camera | Falcon 3EC |  |
| Magnification | × 120,000 |  |
| Total electron dose (e<sup>-</sup>/Å<sup>2</sup>) | 40.0 |  |
| Defocus range (μm) | -0.5 and -2.5 μm |  |
| Pixel size (Å) | 0.889 |  |
| Micrographs (no.) | 5,450 |  |
| Symmetry imposed | C1 | C1 |
| Initial particle images (no.) | 4'980'852 |  |
| Final particle images (no.) | 76'723 | 94,430 |
| Resolution (Å)<br>(FSC threshold) | 3.29<br>(0.143) | 3.52<br>(0.143) |
| Sharpening B-factor (Å<sup>2</sup>) | -115.4 | -127.7 |
| EMDB code |  | 15065 |
| Model refinement |  |  |
| Protein residues | 1862 | 1875 |
| RNA nucleotides | 0 | 6 |
| PLP molecules | 0 | 0 |
| Root-mean-square deviations (RMSDs) |  |  |
| Bond lengths (Å) | 0.002 | 0.002 |
| Bond angles (°) | 0.424 | 0.479 |
| Ramachandran plot |  |  |
| Favored (%) | 98.10 | 97.04 |
| Allowed (%) | 1.90 | 2.91 |
| Disallowed (%) | 0.00 | 0.05 |
| Validation |  |  |
| Molprobrity score | 1.31 | 1.54 |
| Clashscore | 5.73 | 6.67 |
| CC (mask) | 0.85 | 0.75 |
| CC (box) | 0.90 | 0.88 |
| CC (peaks) | 0.90 | 0.66 |
| CC (volume) | 0.84 | 0.74 |
| PDB code | 8C6X | 8A11 |

### catRAPID

The interaction between RNA1-50 stretches and SHMT1 was predicted using catRAPID^39^. CatRAPID estimates the binding potential through van der Waals, hydrogen bonding and secondary structure propensities of both protein and RNA sequences, allowing identification of binding partners with high confidence. As an analysis of about half a million of experimentally validated interactions^81^, the algorithm can separate interacting vs. non-interacting pairs with an area under the curve (AUC) receiver operating characteristic (ROC) curve of 0.78 (with false discovery rate (FDR) significantly below 0.25 when the Z-score values are >2). SHMT regions of 100 residues with a 50-residue overlap were employed in the calculations.

### EMSA assays

EMSA assays were used to assess the binding affinity of different RNAs towards SHMT1, and the ability of the mutants to bind RNA. These experiments were conducted by incubating a fixed amount of RNA with either fixed or increasing concentrations of purified SHMT1 (or variants).

All the components were incubated at 25°C for 30 minutes in 12 μl of binding buffer (20 mM HEPES pH 7.4, 100 mM NaCl) containing 20 μg/ml bovine serum albumin (BSA) and 8% (v/v) glycerol. The reaction mixtures were then subjected to electrophoresis under native conditions using a nondenaturing 4% polyacrylamide gel in 0.5× TBE buffer (45 mM Tris-Borate, 1 mM ethylenediaminetetraacetic acid pH 8.6). For the visualization, gels were stained with SYBR Safe (Invitrogen) in 30 ml of 0.5× TBE and images were acquired using Chemidoc MP Imaging System (Bio-Rad). Densitometric measurements of the free RNA bands were transformed into percentages, the amount of bound RNA (% bound) was then calculated by subtracting the percentage of free RNA from the total, the result was plotted as a function of protein concentration. The apparent dissociation constant (KD) was estimated fitting the data with Equation 1, in which Bmax corresponds to the maximum binding (100%) and [P] corresponds to the concentration of SHMT1 in the reaction mixture.

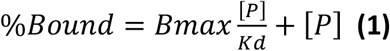

### Activity assays

All assays were carried out at 30 °C in 20 mM K-phosphate buffer at pH 7.2, treated with diethyl pyrocarbonate (DEPC). The serine cleavage reaction was measured using 0.2 μM of purified SHMT1 with fixed 10 mM L-serine and 50 μM THF as substrates while varying the RNA concentration. A spectrophotometric coupled assay, in which the 5,10-methylene-THF produced by the reaction was oxidized by the NAD-dependent *E. coli* 5,10-methylene-THF dehydrogenase was used ^82^.

All obtained inhibition curves were fitted to the equation below, in which [I] corresponds to the RNA concentration:

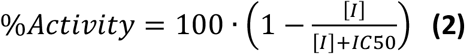

The inhibition kinetics of the SHMT1 by UTR2_1-50_ RNA was carried out as described in Guiducci et al., ^15^ using UTR2_1-50_ RNA in the place of tRNA. The single saturation curves obtained keeping L-serine concentration constant at 10 mM and varying THF (between 3 and 485 μM) at various UTR2_1-50_ RNA concentrations (0, 0.24 e 1.2 μM) were fitted to a modified Michaelis-Menten equation that accounts for substrate inhibition.

All spectrophotometric measurements were performed using a Hewlett-Packard 8453 diode-array spectrophotometer. Fitting of data to equations was carried out with the PRISM software (GraphPad, La Jolla, CA, USA).

### Spectral measurements of SHMT1-amino acid complexes

All SHMT1-amino acid spectra were recorded at 25°C in 50mM Hepes buffer, 150mM NaCl, 5% glycerol at pH 8, in the 260-600 nm range, using Jasco V-650 spectrophotometer. After 30 minutes of incubation at 25°C of UTR2_1-50_ RNA 1:4 SHMT1 18 μM, final concentrations of 10mM serine or glycine respectively were added to the cuvette with 1 cm path length. For the negative control, water with the same volume of RNA solution was added. Acquired spectra were normalized with the respective baseline (SHMT1 buffer) with IgorPro 8 software (Wavemetrics, Lake Oswego, OR, USA) and the differential spectra in the PLP absorption region (315-500 nm) for each experiment were calculated with PRISM software (GraphPad, La Jolla, CA, USA).

### Differential Scanning Fluorimetry (DSF)

DSF assays were performed on a Real Time PCR Instrument (CFX Connect Real Time PCR system, Bio-Rad) using 96-well PCR plates. In each well 1 μM SHMT1 in 20 mM K-phosphate buffer, pH 7.2, and Sypro Orange (2.5x, Thermo Scientific) were mixed in a total volume of 25 μL with various ligands (CHO-THF-5Glu, CHOTHF, lometrexol, tRNA, UTR2_1-50_) in a 96-well PCR plate. Fluorescence was measured from 25 °C to 95 °C in 0.4 °C/30 sec steps (excitation 450-490 nm; detection 560-580 nm). All samples were run in triplicate. Denaturation profiles were analyzed using Graph Pad software after removal of points representing quenching of the fluorescent signal due to postpeak aggregation of protein:dye complexes. All curves were normalized and fitted to a sigmoidal equation to obtain the melting temperatures (Tm). Alternatively, Tm values were obtained by plotting the first derivative of the fluorescence emission as a function of temperature (-dF/dT) by using the Bio-Rad CFX manager software.

### Phylogenetic Analysis

We assessed the presence of the flap motif and, at the same time, of the three major residues responsible for the tetramerization in human SHMT1, i.e., His 135, Arg 137, and Glu 168^18,32^ by modeling SHMT structures from a total of 25 representative species (Figures 6 and S11; Table S2). All the crystal structures and AlphaFold2 models obtained were compared to the crystal structures of SHMT from *E. coli* (PDB 1DFO^83^), and from *H. sapiens* (PDB 1BJ4^32^).

The evolutionary analysis was inferred by using the Maximum Likelihood method and JTT matrix-based model^66^. The tree with the highest log likelihood is shown in Figure S10. Initial tree(s) Evolutionary analyses were conducted in MEGA11^67,68^.

## ACKNOWLEDGMENTS

This work has been supported by funding from Associazione Italiana Ricerca sul Cancro (AIRC) IG2019 n. 23125 (to F.C.); Ministry of Health grant RF-2019-12368888 (to F.C.);LazioInnova, grant A0375-2020-36544 (to F.C. and A.T.); European Union – NextGenerationEU” DD. 3175/2021 E DD. 3138/2021 CN_3: National Center for Gene Therapy and Drugs based on RNA Technology- Project CN 00000041. (to F.C.); Funds from Sapienza University are also gratefully acknowledged (RM11916B46D48441 to F.C., RM12117A5BDD4AEC to G.G., RP12017275CED09F to A.Paiardini., RP12117A675EDA46 and RP1221816B26A29B to A.Paone., RM120172A7AD98EB to S.R., AR2221816C6DB6AE to S.S.); the Pediatric Research Center, Romeo & Enrica Invernizzi Foundation; University of Milano; Associazione Italiana Ricerca sul Cancro (AIRC) grant number MFAG 20447 (A.Paiardini.); MIUR - Ministero dell’Istruzione, dell’Università e della Ricerca (Ministry of Education, University and Research), National Project FSE/FESR-PON “Ricerca e Innovazione” 2014-2020, grant number AIM1887574, CUP E18H19000350007 (M.A.); MIUR-PRIN 2020, grant number 2020PKLEPN_LS3 (L.C.M.).

## AUTHOR CONTRIBUTIONS

F.C. and G.G. conceived and supervised the research. S.S. and F.D.F purified the protein. M.A. helped with the preliminary negative staining analysis. S.S and C.M. prepared the cryo-EM samples and optimized cryo-EM data collection. C.M. processed cryo-EM data. C.M., A.CS, M.B., P.S., G.G, S.S., L.C.M, analyzed the cryo-EM data. G.G. and S.S. interpreted the structure and built the atomic model. A.J.J, A.B performed the Local scale map sharpening. S.S., F.D.F performed the *in vitro* binding assays under the supervision of S.R. and A.Paone. A.T., A.Parroni and R.C. performed the *in vitro* activity assays. A.T., A.Parroni. and S.S performed the DSF analysis. F.D.F and F.R.L. performed the Spectral measurements of SHMT1-amino acid complexes. G.G.T. performed the catRAPID analysis. A.Paiardini. and S.S. performed the Phylogenetic Analysis. G.B. helped in setting up the experiments. All authors discussed the results. F.C., G.G. and S.S. wrote the manuscript. All authors commented on the final version.

## DECLARATION OF INTERESTS

The authors declare no competing interests.

## Notes

**FUNDS**, Associazione Italiana Ricerca sul Cancro (AIRC) IG2019 n. 23125 (to F.C.). Ministry of Health, grant RF-2019-12368888 (to F.C.). LazioInnova, grant A0375-2020-36544 (to F.C. and A.T.). European Union – NextGenerationEU” DD. 3175/2021 E DD. 3138/2021 CN_3: National Center for Gene Therapy and Drugs based on RNA Technology- Project CN 00000041 (to F.C.). Funds from Sapienza University are also gratefully acknowledged (RM11916B46D48441 to F.C., RM12117A5BDD4AEC to G.G., RP12017275CED09F to A.Paiardini., RP12117A675EDA46 and RP1221816B26A29B to A.Paone., RM120172A7AD98EB to S.R., AR2221816C6DB6AE to S.S.). This work has been supported by funding from the Pediatric Research Center, Romeo & Enrica Invernizzi Foundation, University of Milano. Associazione Italiana Ricerca sul Cancro (AIRC) grant number MFAG 20447 (A.Paiardini.). MIUR - Ministero dell’Istruzione, dell’Universitàs e della Ricerca (Ministry of Education, University and Research),National Project FSE/FESR-PON “Ricerca e Innovazione” 2014-2020, grant number AIM1887574, CUP E18H19000350007 (M.A.). MIUR-PRIN 2020, grant number 2020PKLEPN_LS3 (L.C.M.).

### Competing Interest Statement

The authors have declared no competing interest.

## REFERENCES

1. Corley, M., Burns, M.C., and Yeo, G.W. How RNA binding proteins interact with RNA: molecules and mechanisms. 10.1016/j.molcel.2020.03.011.

2. Hentze, M.W., Castello, A., Schwarzl, T., and Preiss, T. (2018). A brave new world of RNA-binding proteins. 10.1038/nrm.2017.130.

3. Li, X., Kazan, H., Lipshitz, H.D., and Morris, Q.D. (2014). Finding the target sites of RNA-binding proteins. Wiley Interdiscip. Rev. RNA 5, 111–130. 10.1002/WRNA.1201.

4. Biamonti, G., and Riva, S. (1994). New insights into the auxiliary domains of eukaryotic RNA binding proteins. FEBS Lett. 340. 10.1016/0014-5793(94)80162-2.

5. Nagy, E., and Rigby, W.F.C. (1995). Glyceraldehyde-3-phosphate Dehydrogenase Selectively Binds AU-rich RNA in the NAD+-binding Region (Rossmann Fold) (*). J. Biol. Chem. 270, 2755–2763. 10.1074/JBC.270.6.2755.

6. Castello, A., Hentze, M.W., and Preiss, T. (2015). Metabolic Enzymes Enjoying New Partnerships as RNA-Binding Proteins. Trends Endocrinol. Metab. 26, 746–757. 10.1016/j.tem.2015.09.012.

7. Volz, K. (2008). The functional duality of iron regulatory protein 1. Curr. Opin. Struct. Biol. 18, 106. 10.1016/J.SBI.2007.12.010.

8. Millet, P., Vachharajani, V., McPhail, L., Yoza, B., and McCall, C.E. (2016). GAPDH Binding to TNF-α mRNA Contributes to Post-Transcriptional Repression in Monocytes: A Novel Mechanism of Communication between Inflammation and Metabolism. J. Immunol. 196, 2541. 10.4049/JIMMUNOL.1501345.

9. Horos, R., Büscher, M., Kleinendorst, R., Alleaume, A.M., Tarafder, A.K., Schwarzl, T., Dziuba, D., Tischer, C., Zielonka, E.M., Adak, A., et al. (2019). The Small Non-coding Vault RNA1-1 Acts as a Riboregulator of Autophagy. Cell 176, 1054–1067.e12. 10.1016/j.cell.2019.01.030.

10. Zhu, Y., Jin, L., Shi, R., Li, J., Wang, Y., Zhang, L., Liang, C.-Z., Narayana, V.K., De Souza, D.P., Thorne, R.F., et al. (2022). The long noncoding RNA glycoLINC assembles a lower glycolytic metabolon to promote glycolysis. Mol. Cell 82, 542–554.e6. 10.1016/j.molcel.2021.11.017.

11. Kerr, A.G., Wang, Z., Wang, N., Kwok, K.H.M., Jalkanen, J., Ludzki, A., Lecoutre, S., Langin, D., Bergo, M.O., Dahlman, I., et al. (2022). The long noncoding RNA ADIPINT regulates human adipocyte metabolism via pyruvate carboxylase. Nat. Commun. 2022 131 13, 1–16. 10.1038/s41467-022-30620-0.

12. Chu, E., and Allegra, C.J. (1996). The role of thymidylate synthase as an RNA binding protein. BioEssays 18, 191–198. 10.1002/BIES.950180306.

13. Chu, E., Takimoto, C., Voeller, D., Grem, J., and Allegra, C. (1993). Specific binding of human dihydrofolate reductase protein to dihydrofolate reductase messenger RNA in vitro. Biochemistry 32, 4756–4760. 10.1021/BI00069A009.

14. Huppertz, I., Perez-Perri, J.I., Mantas, P., Sekaran, T., Schwarzl, T., Russo, F., Ferring-Appel, D., Koskova, Z., Dimitrova-Paternoga, L., Kafkia, E., et al. (2022). Riboregulation of Enolase 1 activity controls glycolysis and embryonic stem cell differentiation. Mol. Cell 82, 2666–2680.e11. 10.1016/J.MOLCEL.2022.05.019.

15. Guiducci, G., Paone, A., Tramonti, A., Giardina, G., Rinaldo, S., Bouzidi, A., Magnifico, M.C., Marani, M., Menendez, J.A., Fatica, A., et al. (2019). The moonlighting RNA-binding activity of cytosolic serine hydroxymethyltransferase contributes to control compartmentalization of serine metabolism. Nucleic Acids Res. 47, 4240–4254. 10.1093/nar/gkz129.

16. Garrow, T.A., Brenner, A.A., Whitehead, V.M., Chen, X.N., Duncan, R.G., Korenberg, J.R., and Shane, B. (1993). Cloning of human cDNAs encoding mitochondrial and cytosolic serine hydroxymethyltransferases and chromosomal localization. J. Biol. Chem. 268, 11910–11916. 10.1016/S0021-9258(19)50286-1.

17. Paone, A., Marani, M., Fiascarelli, A., Rinaldo, S., Giardina, G., Contestabile, R., Paiardini, A., and Cutruzzolà, F. (2014). SHMT1 knockdown induces apoptosis in lung cancer cells by causing uracil misincorporation. Cell Death Dis. 5, 1–11. 10.1038/cddis.2014.482.

18. Giardina, G., Paone, A., Tramonti, A., Lucchi, R., Marani, M., Magnifico, M.C., Bouzidi, A., Pontecorvi, V., Guiducci, G., Zamparelli, C., et al. (2018). The catalytic activity of serine hydroxymethyltransferase is essential for de novo nuclear dTMP synthesis in lung cancer cells. FEBS J. 285, 3238–3253. 10.1111/febs.14610.

19. Monti, M., Guiducci, G., Paone, A., Rinaldo, S., Giardina, G., Liberati, F.R., Cutruzzolá, F., and Tartaglia, G.G. (2021). Modelling of SHMT1 riboregulation predicts dynamic changes of serine and glycine levels across cellular compartments. Comput. Struct. Biotechnol. J. 19, 3034–3041. 10.1016/J.CSBJ.2021.05.019.

20. Zhang, P., and Yang, Q. (2021). Overexpression of SHMT2 Predicts a Poor Prognosis and Promotes Tumor Cell Growth in Bladder Cancer. Front. Genet 12, 682856. 10.3389/fgene.2021.682856.

21. Jain, M., Nilsson, R., Sharma, S., Madhusudhan, N., Kitami, T., Souza, A.L., Kafri, R., Kirschner, M.W., Clish, C.B., and Mootha, V.K. (2012). Metabolite Profiling Identifies a Key Role for Glycine in Rapid Cancer Cell Proliferation. Science 336, 1040. 10.1126/SCIENCE.1218595.

22. Antonov, A., Agostini, M., Morello, M., Minieri, M., Melino, G., Amelio, I., Antonov, A., Agostini, M., Morello, M., Minieri, M., et al. (2014). Bioinformatics analysis of the serine and glycine pathway in cancer cells. Oncotarget 5, 11004–11013. 10.18632/ONCOTARGET.2668.

23. Amelio, I., Cutruzzolá, F., Antonov, A., Agostini, M., and Melino, G. (2014). Serine and glycine metabolism in cancer. Trends Biochem. Sci. 39, 191–198. 10.1016/J.TIBS.2014.02.004.

24. Gu, J., Li, X., Li, H., Jin, Z., and Jin, J. (2019). MicroRNA-198 inhibits proliferation and induces apoptosis by directly suppressing FGFR1 in gastric cancer. Biosci. Rep. 39. 10.1042/BSR20181258.

25. Paiardini, A., Tramonti, A., Schirch, D., Guiducci, G., di Salvo, M.L., Fiascarelli, A., Giorgi, A., Maras, B., Cutruzzolà, F., and Contestabile, R. (2016). Differential 3-bromopyruvate inhibition of cytosolic and mitochondrial human serine hydroxymethyltransferase isoforms, key enzymes in cancer metabolic reprogramming. Biochim. Biophys. Acta 1864, 1506–1517. 10.1016/J.BBAPAP.2016.08.010.

26. Doletha M. E. Szebenyi, ‡, Xiaowen Liu, §, Irina A. Kriksunov, ‡, Patrick J. Stover, *,§ and, and Thiel‡, D.J. (2000). Structure of a Murine Cytoplasmic Serine Hydroxymethyltransferase Quinonoid Ternary Complex: Evidence for Asymmetric Obligate Dimers†. Biochemistry 39, 13313–13323. 10.1021/BI000635A.

27. Scarsdale, J.N., Kazanina, G., Radaev, S., Schirch, V., and Wright, H.T. (1999). Crystal Structure of Rabbit Cytosolic Serine Hydroxymethyltransferase at 2.8 Å Resolution: Mechanistic Implications †, ‡. 10.1021/bi9904151.

28. Zanetti, K.A., and Stover, P.J. (2003). Pyridoxal Phosphate Inhibits Dynamic Subunit Interchange among Serine Hydroxymethyltransferase Tetramers. J. Biol. Chem. 278, 10142–10149. 10.1074/JBC.M211569200.

29. Schirch, V., Shostak, K., Zamora, M., and Guatam-Basak, M. (1991). The origin of reaction specificity in serine hydroxymethyltransferase. J. Biol. Chem. 266, 759–764. https://doi.org/10.1016/S0021-9258(17)35237-7.

30. Stover, P., and Schirch, V. (1992). Enzymatic Mechanism for the Hydrolysis of 5, 10-Methenyltetrahydropteroylglutamate to 5-Formyltetrahydropteroylglutamate by Serine Hydroxymethyltransferase. Biochemistry 31, 2155–2164. 10.1021/bi00122a037.

31. Huang, T., Wang, C., Maras, B., Barra, D., and Schirch, V. (1998). Thermodynamic analysis of the binding of the polyglutamate chain of 5-formyltetrahydropteroylpolyglutamates to serine hydroxymethyltransferase. Biochemistry 37, 13536–13542. 10.1021/bi980827u.

32. Renwick, S.B., Snell, K., and Baumann, U. (1998). The crystal structure of human cytosolic serine hydroxymethyltransferase: A target for cancer chemotherapy. Structure 6, 1105–1116. 10.1016/S0969-2126(98)00112-9.

33. Szebenyi, D.M.E., Liu, X., Kriksunov, I.A., Stover, P.J., and Thiel, D.J. (2000). Structure of a murine cytoplasmic serine hydroxymethyltransferase quinonoid ternary complex: Evidence for asymmetric obligate dimers. Biochemistry 39, 13313–13323. 10.1021/BI000635A/ASSET/IMAGES/LARGE/BI000635AF00008.JPEG.

34. Schirch, V., and Szebenyi, D.M.E. (2005). Serine hydroxymethyltransferase revisited. Curr. Opin. Chem. Biol. 9, 482–487. 10.1016/j.cbpa.2005.08.017.

35. Ubonprasert, S., Jaroensuk, J., Pornthanakasem, W., Kamonsutthipaijit, N., Wongpituk, P., Mee-udorn, P., Rungrotmongkol, T., Ketchart, O., Chitnumsub, P., Leartsakulpanich, U., et al. (2019). A flap motif in human serine hydroxymethyltransferase is important for structural stabilization, ligand binding, and control of product release. J. Biol. Chem. 294, 10490–10502. 10.1074/JBC.RA119.007454.

36. Chu, E., and Allegra, C.J. (1996). The role of thymidylate synthase as an RNA binding protein. BioEssays 18, 191–198. 10.1002/BIES.950180306.

37. A. Ercikan-Abali, E., Banerjee, D., C. Waltham, M., Skacel, N., W. Scotto, K., and R. Bertino, J. (1997). Dihydrofolate Reductase Protein Inhibits Its Own Translation by Binding to Dihydrofolate Reductase mRNA Sequences within the Coding Region. Biochemistry 36, 12317–12322. 10.1021/bi971026e.

38. Cieśla, J. (2006). Metabolic enzymes that bind RNA: Yet another level of cellular regulatory network? Acta Biochim. Pol. 53, 11–32. 10.18388/abp.2006_3360.

39. Bellucci, M., Agostini, F., Masin, M., and Tartaglia, G.G. (2011). Predicting protein associations with long noncoding RNAs. Nat. Methods 8, 444–445. 10.1038/NMETH.1611.

40. Fu, T.-F., Scarsdale, J.N., Kazanina, G., Schirch, V., and Wright, H.T. (2003). Location of the Pteroylpolyglutamate-binding Site on Rabbit Cytosolic Serine Hydroxymethyltransferase *. J. Biol. Chem. 278, 2645–2653. 10.1074/JBC.M210649200.

41. Stover, P.J., and Field, M.S. (2011). Trafficking of Intracellular Folates. Adv. Nutr. 2, 325–331. 10.3945/AN.111.000596.

42. Zeng, H., Chen, Z.-S., Belinsky, M.G., Rea, P.A., and Kruh, G.D. (2001). Transport of Methotrexate (MTX) and Folates by Multidrug Resistance Protein (MRP) 3 and MRP1: Effect of Polyglutamylation on MTX Transport 1. CANCER Res. 61, 7225–7232.

43. Cornish-Bowden, A. (1995). Fundamentals of enzyme kinetics, Revised Edition. (Portland Press Ltd., London, UK.).

44. Monod, J., Changeux, J.P., and Jacob, F. (1963). Allosteric proteins and cellular control systems. J. Mol. Biol. 6, 306–329. 10.1016/S0022-2836(63)80091-1.

45. Barlow, J.N., Conrath, K., and Steyaert, J. (2009). Substrate-dependent modulation of enzyme activity by allosteric effector antibodies. Biochim. Biophys. Acta - Proteins Proteomics 1794, 1259–1268. 10.1016/J.BBAPAP.2009.03.019.

46. Amornwatcharapong, W., Maenpuen, S., Chitnumsub, P., Leartsakulpanich, U., and Chaiyen, P. (2017). Human and Plasmodium serine hydroxymethyltransferases differ in rate-limiting steps and pH-dependent substrate inhibition behavior. Arch. Biochem. Biophys. 630, 91–100. 10.1016/J.ABB.2017.07.017.

47. Schirch, V., Shostak, K., Zamora, M., and Gautam-Basak, M. (1991). The origin of reaction specificity in serine hydroxymethyltransferase. J. Biol. Chem. 266, 759–764. 10.1016/s0021-9258(17)35237-7.

48. Cobbold, S.A., Tutor, M. V, Frasse, P., McHugh, E., Karnthaler, M., Creek, D.J., John, A.O., Tilley, L., Ralph, S.A., and McConville, M.J. (2021). Non-canonical metabolic pathways in the malaria parasite detected by isotope-tracing metabolomics. Mol. Syst. Biol. 17, e10023. 10.15252/MSB.202010023.

49. Ducker, G., Chen, L., Morscher, R., Ghergurovich, J., Esposito, M., Teng, X., Kang, Y., and Rabinowitz, J. (2016). Reversal of Cytosolic One-Carbon Flux Compensates for Loss of the Mitochondrial Folate Pathway. Cell Metab. 23, 1140–1153. 10.1016/J.CMET.2016.04.016.

50. Minton, D.R., Nam, M., McLaughlin, D.J., Shin, J., Bayraktar, E.C., Alvarez, S.W., Sviderskiy, V.O., Papagiannakopoulos, T., Sabatini, D.M., Birsoy, K., et al. (2018). Serine Catabolism by SHMT2 Is Required for Proper Mitochondrial Translation Initiation and Maintenance of Formylmethionyl-tRNAs. Mol. Cell 69, 610–621.e5. 10.1016/J.MOLCEL.2018.01.024.

51. Morscher, R.J., Ducker, G.S., Li, S.H.-J., Mayer, J.A., Gitai, Z., Sperl, W., and Rabinowitz, J.D. (2018). Mitochondrial translation requires folate-dependent tRNA methylation. Nat. Publ. Gr. 10.1038/nature25460.

52. Hanahan, D., and Weinberg, R.A. (2011). Hallmarks of Cancer: The Next Generation. Cell 144, 646–674. 10.1016/j.cell.2011.02.013.

53. Vicens, Q., and Kieft, J.S. (2022). Thoughts on how to think (and talk) about RNA structure. Proc. Natl. Acad. Sci. U. S. A. 119, e2112677119. 10.1073/PNAS.2112677119/SUPPL_FILE/PNAS.2112677119.SAPP.PDF.

54. Ganser, L.R., Kelly, M.L., Herschlag, D., and Al-Hashimi, H.M. (2019). The roles of structural dynamics in the cellular functions of RNAs. Nat. Rev. Mol. Cell Biol. 20, 474. 10.1038/S41580-019-0136-0.

55. Ralser, M. (2014). The RNA world and the origin of metabolic enzymes. Biochem. Soc. Trans. 42, 985–988. 10.1042/BST20140132.

56. Kirschning, A. (2021). Coenzymes and Their Role in the Evolution of Life. Angew. Chemie Int. Ed. 60, 6242–6269. 10.1002/ANIE.201914786.

57. Giacobelli, V.G., Fujishima, K., Lepšík, M., Tretyachenko, V., Kadavá, T., Makarov, M., Bednárová, L., Novák, P., and Hlouchová, K. (2022). In Vitro Evolution Reveals Noncationic Protein–RNA Interaction Mediated by Metal Ions. Mol. Biol. Evol. 39. 10.1093/MOLBEV/MSAC032.

58. Tramonti, A., Nardella, C., Salvo, M.L. di, Barile, A., Cutruzzolà, F., and Contestabile, R. (2018). Human Cytosolic and Mitochondrial Serine Hydroxymethyltransferase Isoforms in Comparison: Full Kinetic Characterization and Substrate Inhibition Properties. Biochemistry 57, 6984–6996. 10.1021/ACS.BIOCHEM.8B01074.

59. Wong, Y., and Rosindell, J. (2022). Dynamic visualisation of million-tip trees: The OneZoom project. Methods Ecol. Evol. 13, 303–313. 10.1111/2041-210X.13766.

60. Spizzichino, S., Pampalone, G., Dindo, M., Bruno, A., Romani, L., Cutruzzolá, F., Zelante, T., Pieroni, M., Cellini, B., and Giardina, G. (2021). Crystal structure of Aspergillus fumigatus AroH, an aromatic amino acid aminotransferase. Proteins. 10.1002/prot.26234.

61. Giardina, G., Montioli, R., Gianni, S., Cellini, B., Paiardini, A., Voltattorni, C.B., and Cutruzzolà, F. (2011). Open conformation of human DOPA decarboxylase reveals the mechanism of PLP addition to Group II decarboxylases. Proc. Natl. Acad. Sci. U. S. A. 108, 20514. 10.1073/PNAS.1111456108.

62. Dindo, M., Pascarelli, S., Chiasserini, D., Grottelli, S., Costantini, C., Uechi, G.I., Giardina, G., Laurino, P., and Cellini, B. (2022). Structural dynamics shape the fitness window of alanine:glyoxylate aminotransferase. Protein Sci. 31, e4303. 10.1002/PRO.4303.

63. Tiwari, S.P., Fuglebakk, E., Hollup, S.M., Skjærven, L., Cragnolini, T., Grindhaug, S.H., Tekle, K.M., and Reuter, N. (2014). WEBnmat v2.0: Web server and services for comparing protein flexibility. BMC Bioinformatics 15, 1–12. 10.1186/S12859-014-0427-6/FIGURES/4.

64. Di Salvo, M.L., Scarsdale, J.N., Kazanina, G., Contestabile, R., Schirch, V., and Wright, H.T. (2013). Structure-Based Mechanism for Early PLP-Mediated Steps of Rabbit Cytosolic Serine Hydroxymethyltransferase Reaction. Biomed Res. Int. 2013, 13. 10.1155/2013/458571.

65. Oliveira, E.F., Cerqueira, N.M.F.S.A., Fernandes, P.A., and Ramos, M.J. (2011). Mechanism of Formation of the Internal Aldimine in Pyridoxal 5 0-Phosphate-Dependent Enzymes. J. Am. Chem. Soc 133, 15496–15505. 10.1021/ja204229m.

66. Jones, D.T., Taylor, W.R., and Thornton, J.M. (1992). The rapid generation of mutation data matrices from protein sequences. Bioinformatics 8, 275–282. 10.1093/bioinformatics/8.3.275.

67. Tamura, K., Stecher, G., and Kumar, S. (2021). MEGA11: Molecular Evolutionary Genetics Analysis Version 11. Mol. Biol. Evol. 38, 3022–3027. 10.1093/molbev/msab120.

68. Stecher, G., Tamura, K., and Kumar, S. (2020). Molecular Evolutionary Genetics Analysis (MEGA) for macOS. 10.1093/molbev/msz312.

69. Spizzichino, S., Boi, D., Boumis, G., Lucchi, R., Liberati, F.R., Capelli, D., Montanari, R., Pochetti, G., Piacentini, R., Parisi, G., et al. (2021). Cytosolic localization and in vitro assembly of human de novo thymidylate synthesis complex. FEBS J. 10.1111/FEBS.16248.

70. Zivanov, J., Nakane, T., Forsberg, B.O., Kimanius, D., Hagen, W.J.H., Lindahl, E., and Scheres, S.H.W. (2018). New tools for automated high-resolution cryo-EM structure determination in RELION-3. Elife 7. 10.7554/ELIFE.42166.

71. Rohou, A., and Grigorieff, N. (2015). CTFFIND4: Fast and accurate defocus estimation from electron micrographs. J. Struct. Biol. 192, 216–221. 10.1016/J.JSB.2015.08.008.

72. Rosenthal, P.B., and Henderson, R. (2003). Optimal determination of particle orientation, absolute hand, and contrast loss in single-particle electron cryomicroscopy. J. Mol. Biol. 333, 721–745. 10.1016/j.jmb.2003.07.013.

73. Jakobi, A.J., Wilmanns, M., and Sachse, C. (2017). Model-based local density sharpening of cryo-EM maps. Elife 6. 10.7554/ELIFE.27131.

74. Bharadwaj, A., and Jakobi, A.J. (2022). Electron scattering properties of biological macromolecules and their use for cryo-EM map sharpening. Faraday Discuss. 240, 168–183. 10.1039/D2FD00078D.

75. Jakobi, A.J., Wilmanns, M., and Sachse, C. (2017). Model-based local density sharpening of cryo-EM maps. Elife 6. 10.7554/ELIFE.27131.

76. Pettersen, E.F., Goddard, T.D., Huang, C.C., Couch, G.S., Greenblatt, D.M., Meng, E.C., and Ferrin, T.E. (2004). UCSF Chimera—A visualization system for exploratory research and analysis. J. Comput. Chem. 25, 1605–1612. 10.1002/JCC.20084.

77. Emsley, P., Lohkamp, B., Scott, W., and Cowtan, K. (2010). Features and development of Coot. Acta Crystallogr. D. Biol. Crystallogr. 66, 486–501. 10.1107/S0907444910007493.

78. Vendeix, F.A.P., Dziergowska, A., Gustilo, E.M., Graham, W.D., Sproat, B., Malkiewicz, A., and Agris, P.F. (2008). Anticodon Domain Modifications Contribute Order to tRNA for Ribosome-Mediated Codon Binding†‡. Biochemistry 47, 6117–6129. 10.1021/BI702356J.

79. Adams, P.D., Afonine, P. V, Bunkó, G., Chen, V.B., Davis, I.W., Echols, N., Headd, J.J., Hung, L.-W., Kapral, G.J., Grosse-Kunstleve, R.W., et al. (2010). Biological Crystallography PHENIX: a comprehensive Python-based system for macromolecular structure solution. Res. Pap. Acta Cryst 66, 213–221. 10.1107/S0907444909052925.

80. Afonine, P.V., Poon, B.K., Read, R.J., Sobolev, O.V., Terwilliger, T.C., Urzhumtsev, A., Adams, P.D., and IUCr (2018). Real-space refinement in PHENIX for cryo-EM and crystallography. urn:issn:2059-7983 74, 531–544. 10.1107/S2059798318006551.

81. Lang, B., Armaos, A., and Tartaglia, G.G. (2019). RNAct: Protein-RNA interaction predictions for model organisms with supporting experimental data. Nucleic Acids Res. 47, D601–D606. 10.1093/NAR/GKY967.

82. Fu, T., Rife, J., and Schirch, V. (2001). The role of serine hydroxymethyltransferase isozymes in one-carbon metabolism in MCF-7 cells as determined by (13)C NMR. Arch. Biochem. Biophys. 393, 42–50. 10.1006/ABBI.2001.2471.

83. Scarsdale, J.N., Radaev, S., Kazanina, G., Schirch, V., and Wright, H.T. (2000). Crystal structure at 2.4 A resolution of E. coli serine hydroxymethyltransferase in complex with glycine substrate and 5-formyl tetrahydrofolate. J. Mol. Biol. 296, 155–168. 10.1006/JMBI.1999.3453.

