## supplementary figures for "THE RIBOREGULATION MECHANISM OF HUMAN SERINE HYDROXYMETHYLTRANSFERASE IS ROOTED IN AN ALLOSTERIC SWITCH"

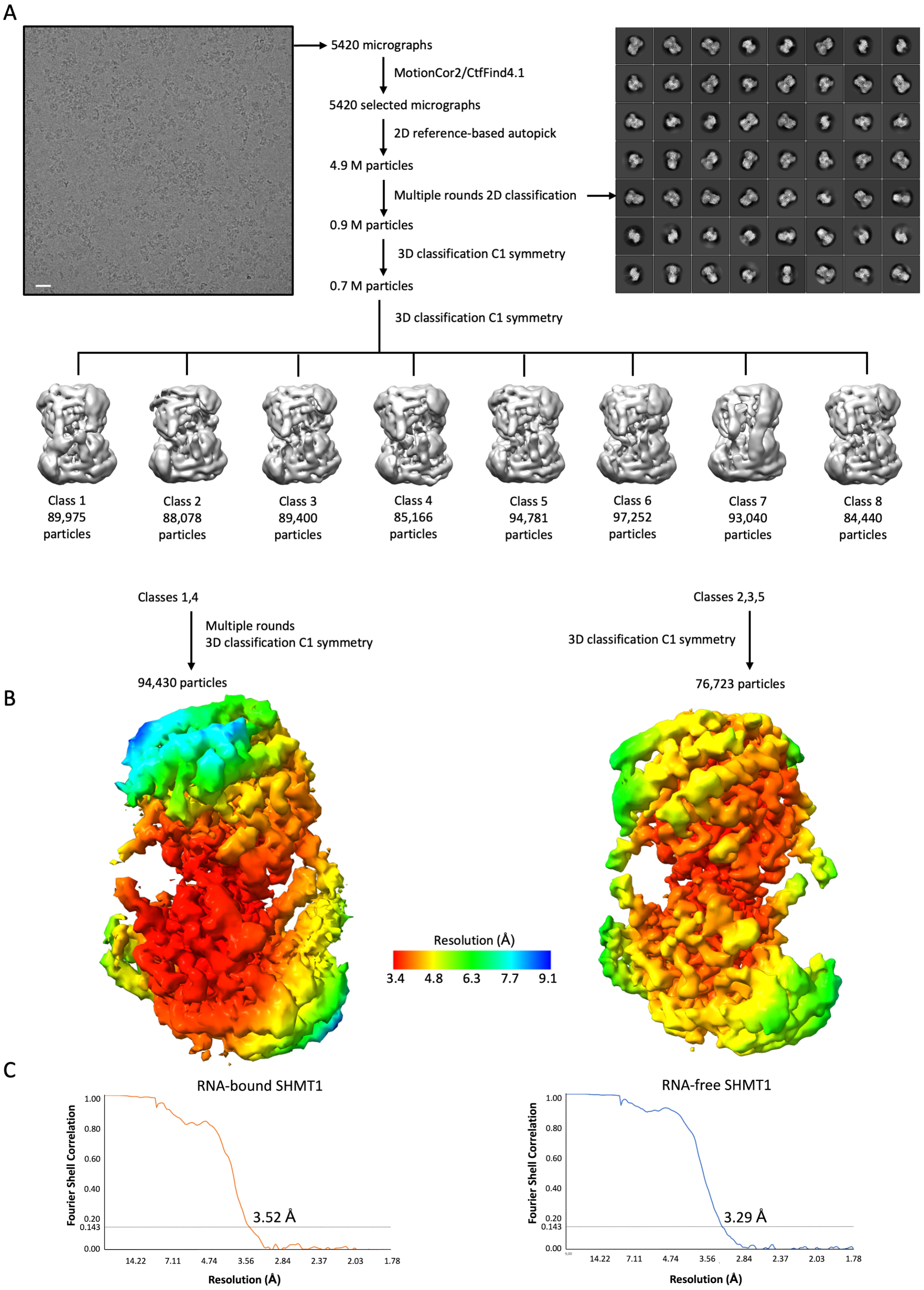


***Figure S1. Cryo-EM workflow. A)*** *Schematic workflow of the image processing and single particle analysis. Top left: representative micrograph of the analysed sample containing 5.7 µM of SHMT1 and 28.5 µM RNA (UTR2 1-50). The scale bar is 50 nm. Top right: All classes from the last 2D classification run. After junk particles had been removed in the first round of 3D classification, the second round revealed 8 different classes, some showing the*

*RNA extra-density and others not. The indicated classes were selected, and two distinct paths were taken to isolate the two most significant SHMT1 states: with and without the RNA-molecule bound.* ***B)*** *The final post-processed volumes are shown for the RNA-bound (left) and the RNA-free (right) SHMT1 structure. Both maps are coloured according to the local resolution, estimated in RELION's LocRes. The colour legend is shown in between the two volumes.* ***C)*** *Gold-standard FSC curves of the SHMT1-RNA complex (left) and the RNA-free enzyme (right). The final resolutions are indicated.*

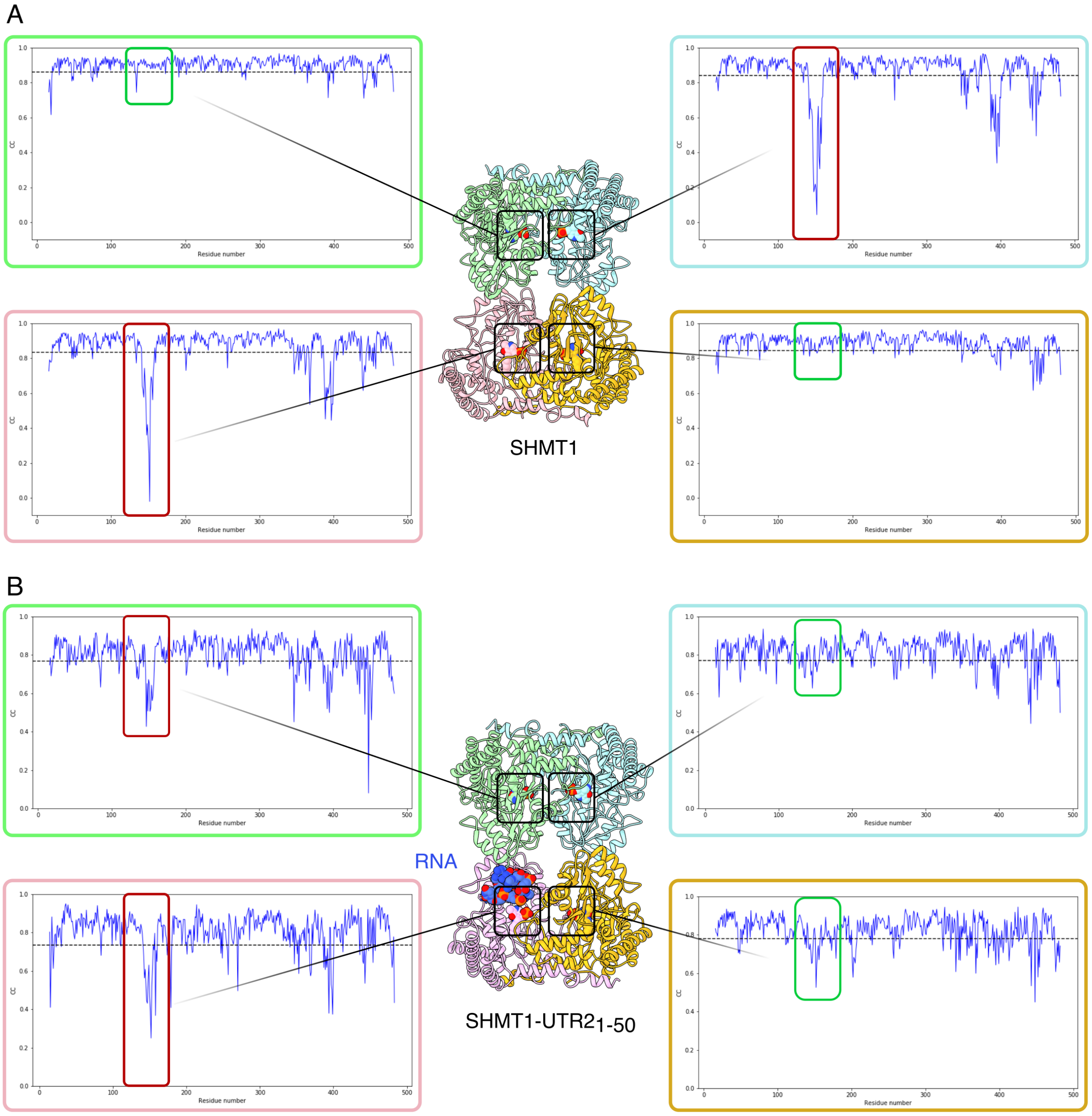


***Figure S2. RNA binding affects the active site conformation.*** *The correlation coefficient plotted against residue number of the RNA-free structure (panel A) or the RNA complex (panel B) shows a marked decrease in the region of the active sites (boxed in red) of the chains in the open conformation, in agreement with the lack of electron*

*density observed in this region. The chains in the open conformation also display an increased flexibility of the C-terminal domain.*

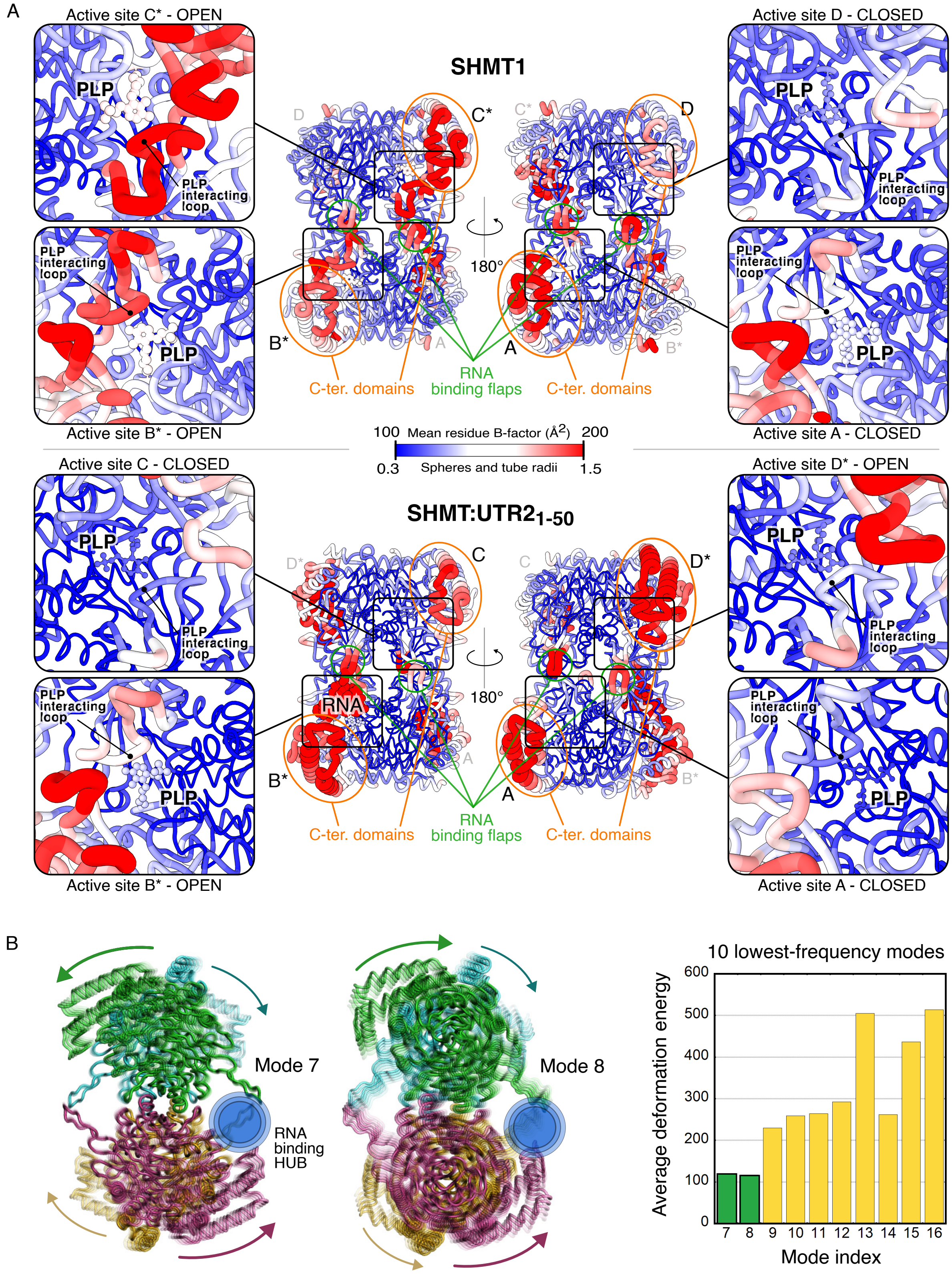


***Figure S3. B-factor mapping and normal mode analysis. A)*** *Cryo-EM structure of SHMT1 and of the SHMT1-RNA complex rendered according to mean B-factors. The same B-factor scale was used for both structures as the*

*mean B-factor is comparable (133 Å$^2$ for SHMT1 and 136 Å$^2$ for the complex). For all chains in both structures the C-terminus domains and the flap are the most dynamic regions, as observed in other PLP-dependent enzymes [60–62]. On the contrary, B-factors of PLP and especially of the region interacting with the cofactor (labelled PLP interacting loop) are different; higher for the chains in the open conformation and lower for those in the closed one.*

***B)*** *The first two lowest energy normal modes are shown together with the plot of the corresponding deformation energies computed using the crystal structure of SHMT1 using WEBnm@ v2.0: (http://apps.cbu.uib.no/webnma3)[63]. The arrows indicate the direction of movement for each subunit.*

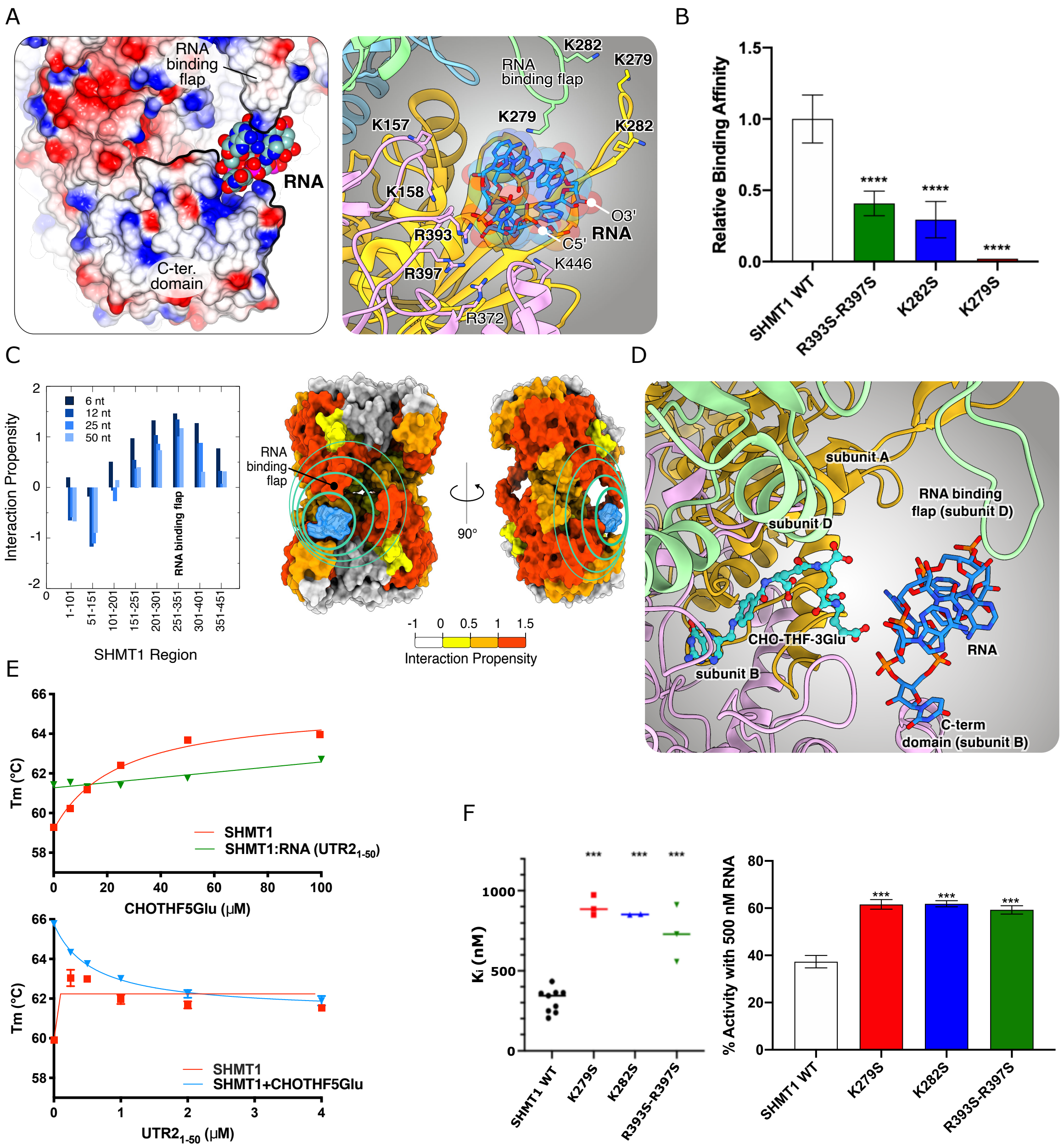


***Figure S4. Comparison between the different RNA oligonucleotides. A)*** *Analysis of the migration of 2 µM RNA (UTR2 1-50 white, UTR2 1-50 Rv orange, UTR2 loop length 8 nt light blue, UTR2 loop length 28 nt pink and dsRNA purple) in the presence of 4- fold excess of SHMT1 by electrophoretic shift assays (EMSA). The analysis shows that the optimal loop length is around 12 nt. The UTR2 1-50 and UTR2 1-50 Rv fragments bearing a loop of this size display the higher affinity towards SHMT1, and were set at 100% and 98%, respectively. When changing the length of the loop the affinity decreases (UTR2 loop length 8 nt has a binding affinity of 77% with respect to UTR2 1-50, while the UTR2 loop length 28 nt of 53%). When using a dsRNA (obtained by annealing the UTR2 1-50 and UTR2 1-50 Rv) the affinity drastically decreases (11%), confirming that the presence of a looped region is required to obtain a good binding affinity. Experiments were run in triplicate (n=3).* ***B)*** *Secondary structures of UTR2 1-50*

*(AUA AAG AAA AAA GCG GUG AGU GGG CGA ACU ACA AUU CCC AAA AGG CCA CA), UTR2 1-50 Rv (UGU GGC CUU UUG GGA AUU GUA GUU CGC CCA CUC ACC GCU UUU UUC UUU AU), UTR2 loop length 8 nt (AUA AAG AAA AAA GCG GUG AGU GGG CGA AUU CCC AAA AGG CCA CA), UTR2 loop lenght 28 nt (AUA AAG AAA AAA GCG GUG AGU GGG CGA ACU AAA AAA AAA AAA ACA AUU CCC AAA AGG CCA CA predicted by using Vienna RNAfold web server: RNAfold web server).* ***C****) EMSA assays using oligonucleotides with similar structure but different sequence***. *The assay was performed by incubating 0.24 µM of either UTR2$_{1-50}$ (AUA AAG AAA AAA GCG GUG AGU GGG CGA ACU ACA AUU CCC AAA AGG CCA CA) or UTR2$_{1-50}$ reverse (rv) (UGU GGC CUU UUG GGA AUU GUA GUU CGC CCA CUC ACC GCU UUU UUC UUU AU) with the indicated concentrations of SHMT1 WT (µM) (n=3).* ***D)*** *Calculation of binding affinity from the EMSA experiments in (C)***.** *The apparent dissociation constants (Kd) of SHMT1-UTR2$_{1-50}$ and SHMT1-UTR2$_{1-50}$rv complexes are respectively 0.27 ± 0.03 µM and 0.21 ± 0.01 µM. These results show that the presence of the hairpin region is crucial for an optimal binding to SHMT1.*

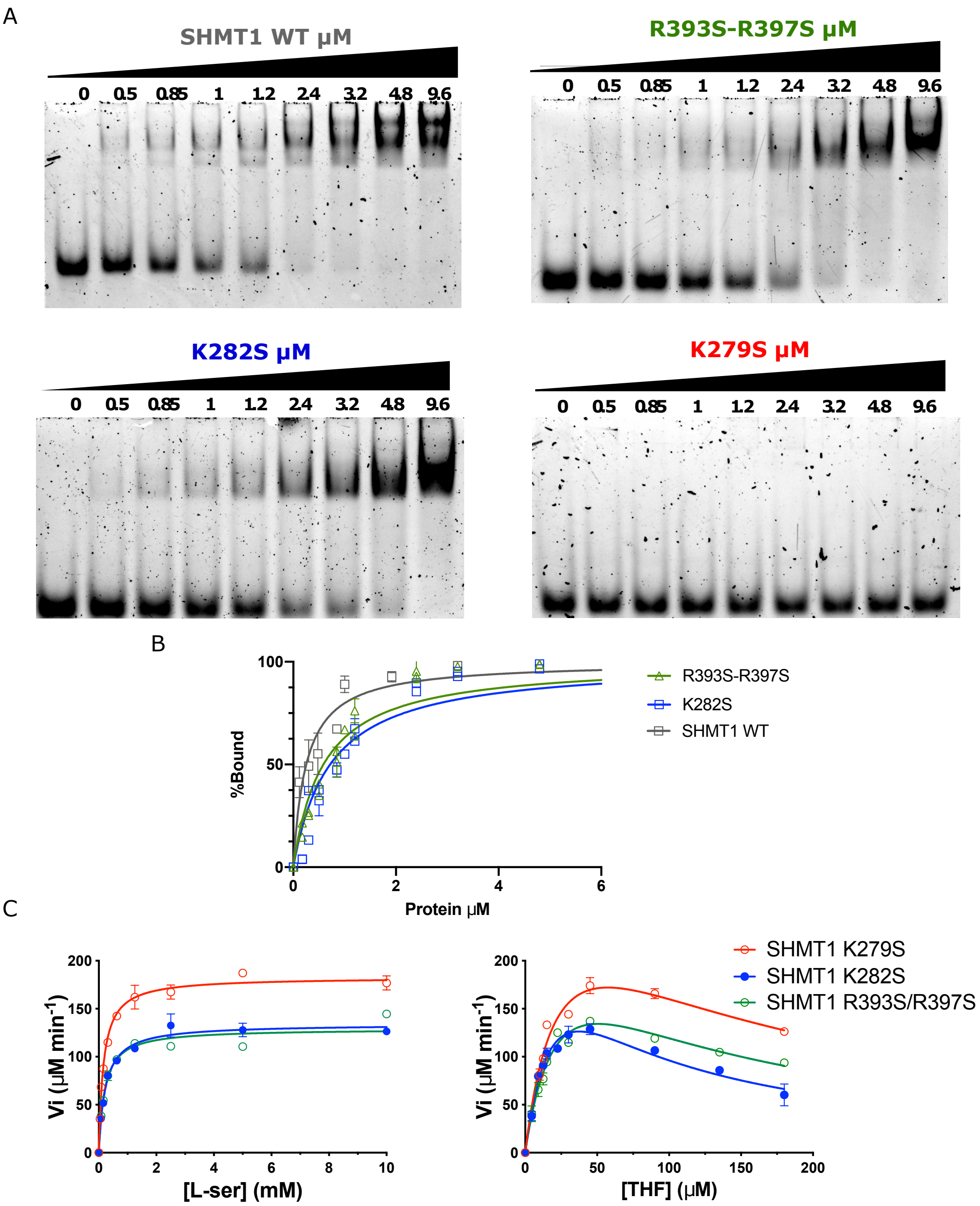


| | $K_M$THF (µM) | $K_i$THF(µM) | $K_M$ser(µM) | $K_{cat}$ ser ($min^{-1}$) |
|---|---|---|---|---|
| SHMT1 WT | 10.2 ± 0.7 | 297 ± 69 | 225 ± 0.01 | 715 |
| K279S | 26.4 ± 7.2 | 125 ± 41 | 163 ± 13 | 689 |
| K282S | 27.7 ± 8.1 | 50 ± 15 | 229 ± 23 | 479 |
| R393S-R397S | 28.4 ± 7.3 | 91 ± 28 | 202 ± 29 | 396 |

***Figure S5. RNA binding propensities and Serine cleavage activity of SHMT1 mutants. A and B)*** *Binding affinity of the SHMT1 variants towards UTR2$_{1-50}$. The apparent dissociation constants (Kd) of SHMT1 mutants and UTR2$_{1-50}$ were calculated from the EMSA experiments. SHMT1 K279S shows the lowest affinity towards RNA binding only 40% of the RNA at the highest protein excess (40-fold, not shown). The Kd of SHMT1 WT is 0.27 ± 0.03 µM,* *the Kd of SHMT1 K282S is 1.24 ± 0.18 µM and that of SHMT1 R393S-R397S is 0.87 ± 0.13 µM, whereas the Kd of SHMT1 K279S cannot be calculated given the low affinity. All the experimental data, acquired in at least three independent experiments, were calculated using equation 1 (see Materials and Methods) and fitted by using a hyperbolic function.* ***C)*** *Kinetic characterization of SHMT1 variants. All the experiments were performed in triplicates.*

A

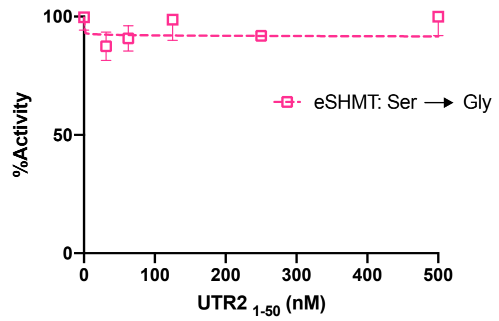


B

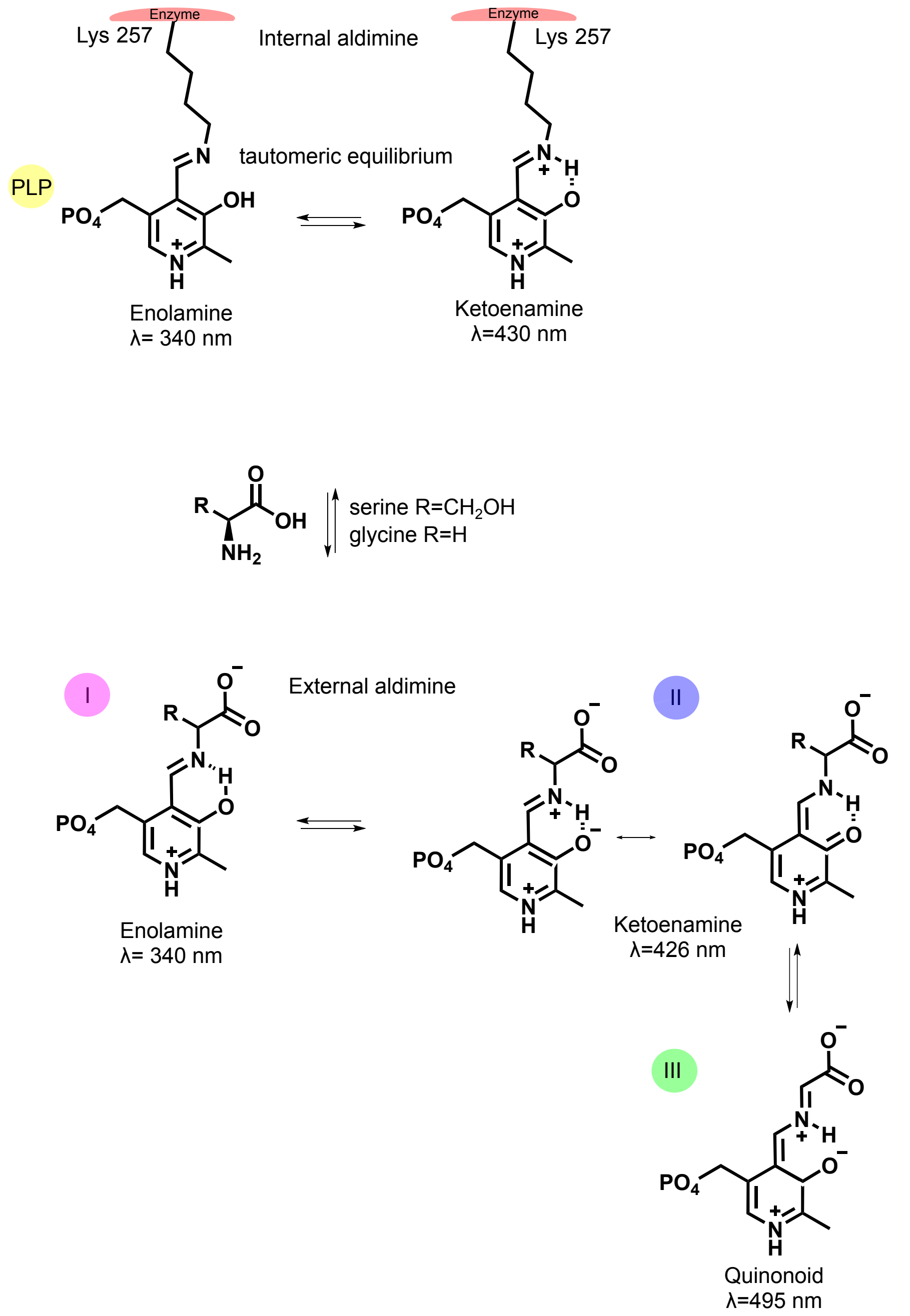


***Figure S6. Bacterial SHMT serine cleavage reactions and proposed reaction mechanism of amino acid binding to SHMT A)*** *Effect of UTR2$_{1-50}$ on bacterial SHMT serine cleavage reactions. The serine cleavage activity of bacterial SHMT (from Escherichia coli, eSHMT) in the presence of UTR2$_{1-50}$ was measured. No effect of the RNA on the bacterial enzyme activity was present in the range of RNA concentration explored (n=3).* ***B)*** *Proposed*

*reaction mechanism of amino acid binding to SHMT [30,32,64,65]. Initially, the PLP cofactor is covalently bound as a Schiff base to the sidechain of Lys257 forming the internal aldimine, which is in a tautomeric equilibrium. When the amino acids are added the internal aldimine is formed. As previously seen in literature [64] and in fig 5A, when glycine is added it is possible to observe all the 3 PLP intermediates (I,II and III). Instead, when serine is added, species II is the main species observed in solution[32].*

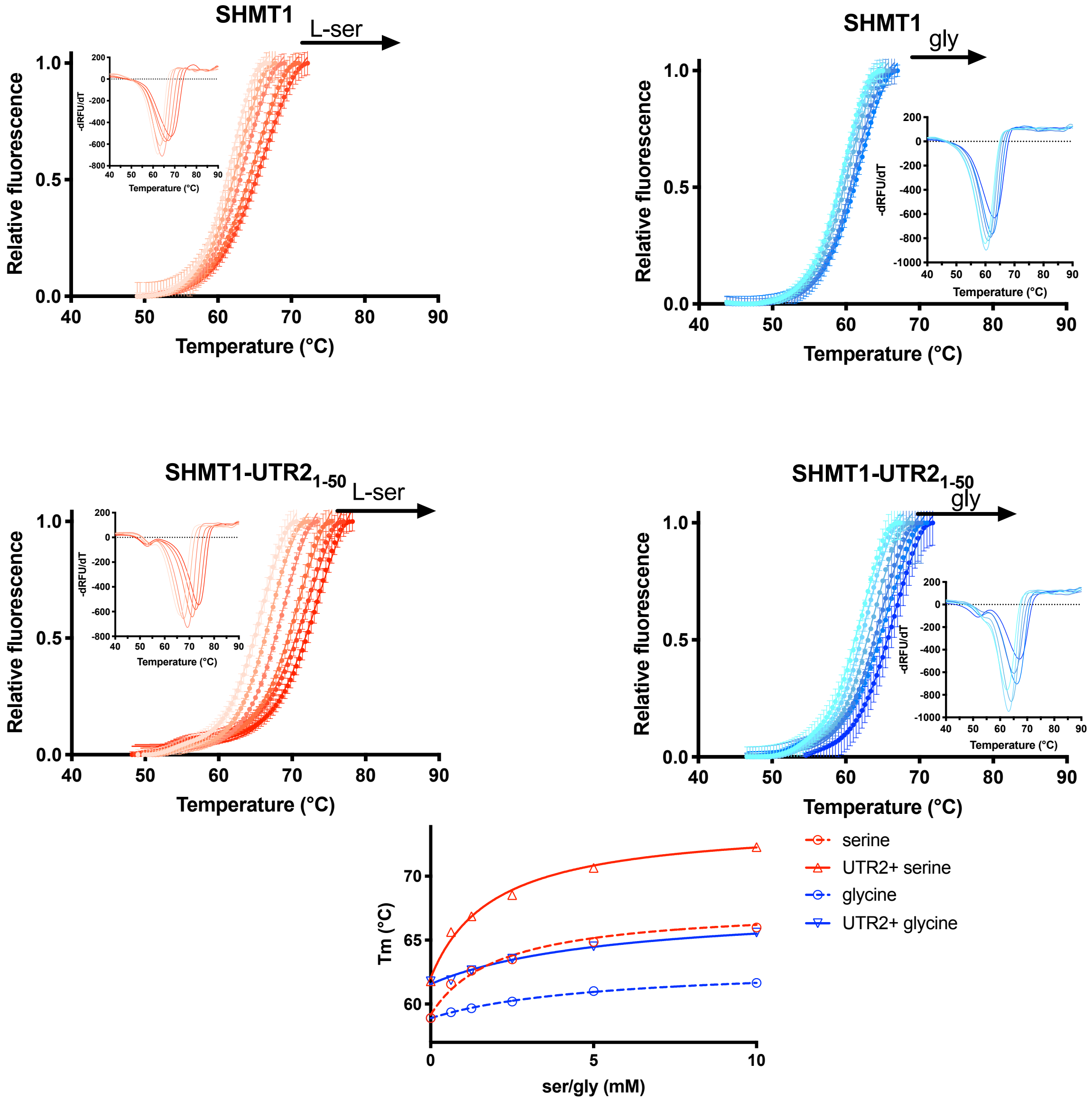


***Figure S7. Effect of amino acid substrates on the stability of SHMT1.*** *DSF experiments probing the effect of serine (red) and glycine (blue) (10 mM) on the stability of the SHMT1 and SHMT1:UTR2$_{1-50}$ complex. 0.5 µM of UTR2$_{1-50}$ were mixed with 2 µM SHMT1, (n=3).*

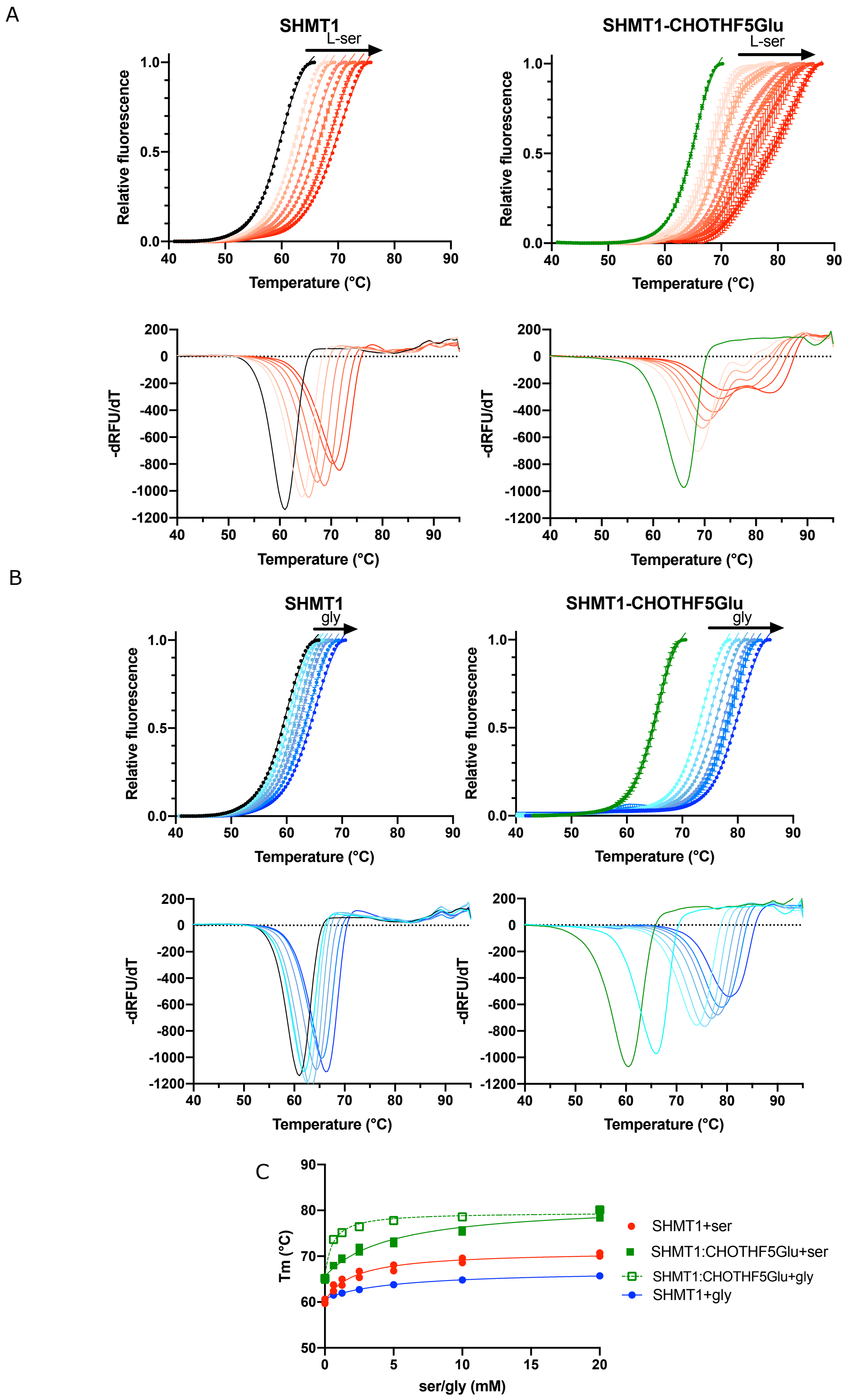


***Figure S8. Titration of SHMT1 (or SHMT1: CHOTHF5Glu) with serine or glycine****. DSF titration with either serine (A) or glycine(B) added to a solution of SHMT1 (2 µM) and 50 µM CHOTHF5Glu (n=3). C) Plot of the observed Tm values as a function of amino acid concentration.*

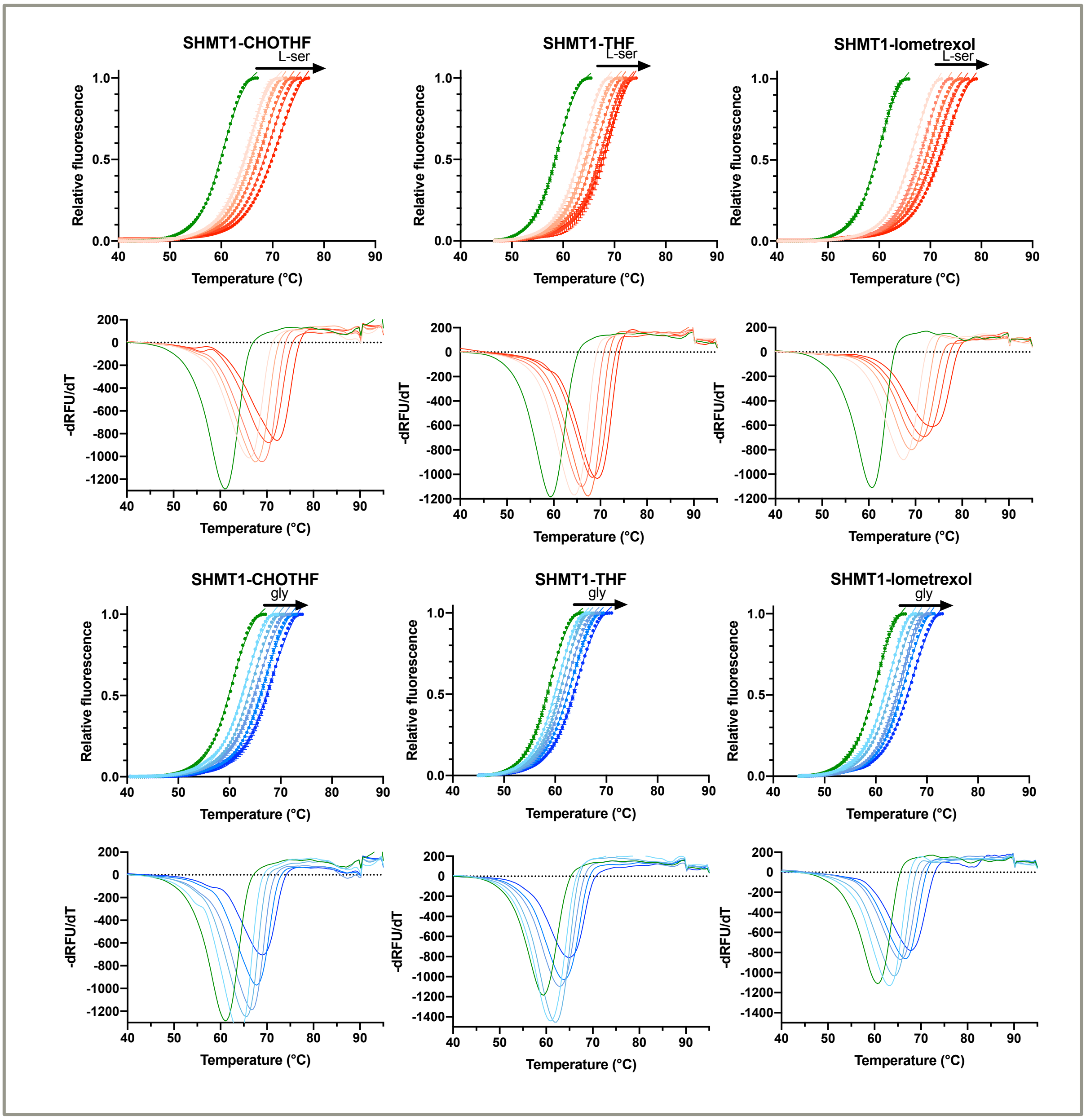


***Figure S9. Titration of SHMT1 (or SHMT1: folates) with serine or glycine.*** *DSF titration with either serine or glycine added to a solution of SHMT1 (2 µM) and 50 µM THF, CHO-THF or lometrexol.*

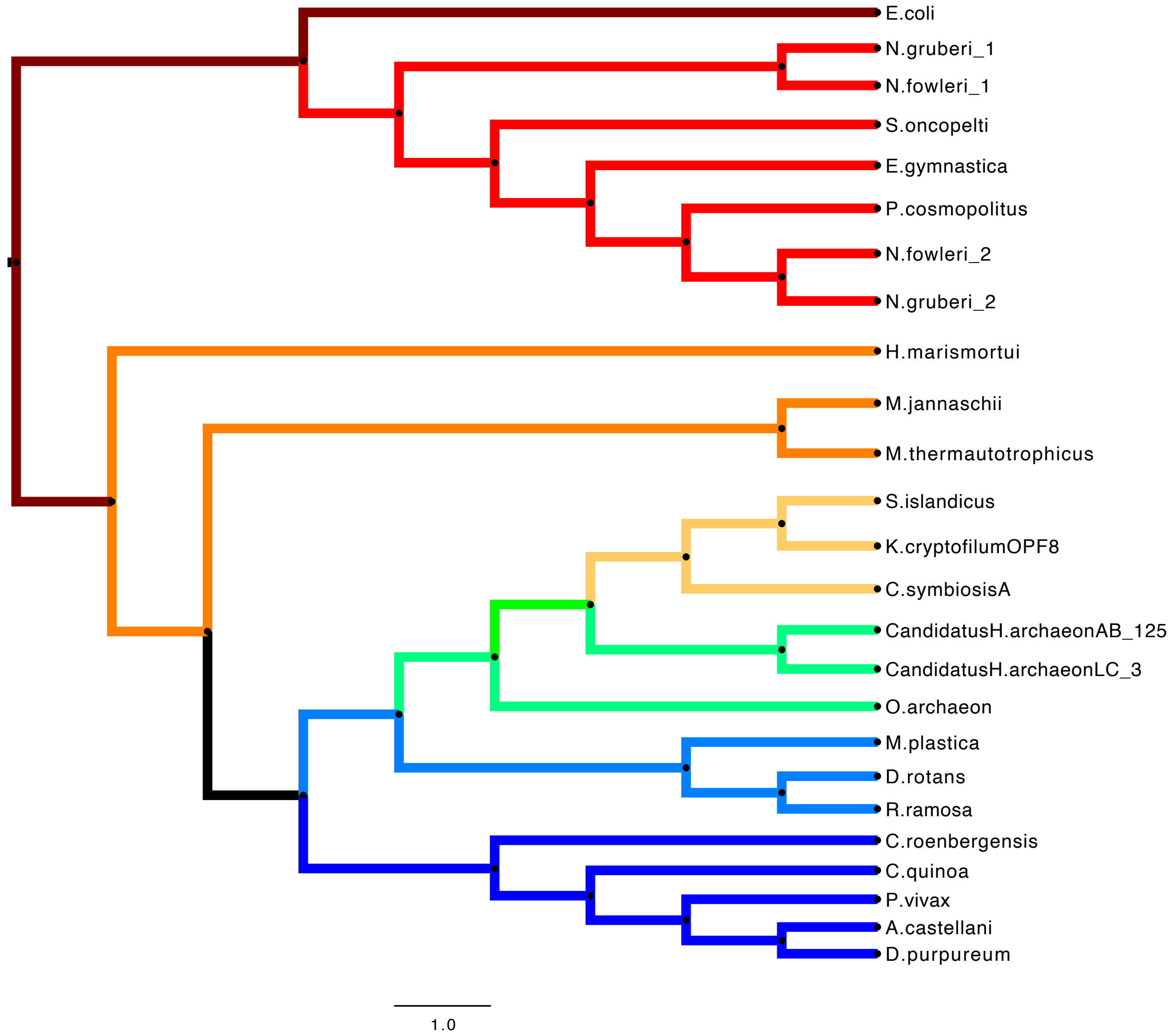


***Figure S10. Evolutionary analysis by Maximum Likelihood method.*** *The evolutionary analysis was inferred by using the Maximum Likelihood method and JTT matrix-based model [66]. The tree with the highest log likelihood is shown. Initial tree(s) for the heuristic search were obtained automatically by applying Neighbor-Join and BioNJ algorithms to a matrix of pairwise distances estimated using the JTT model, and then selecting the topology with superior log likelihood value. This analysis involved 25 amino acid sequences. There was a total of 538 positions in the final dataset. Evolutionary analyses were conducted in MEGA11 [67,68].*

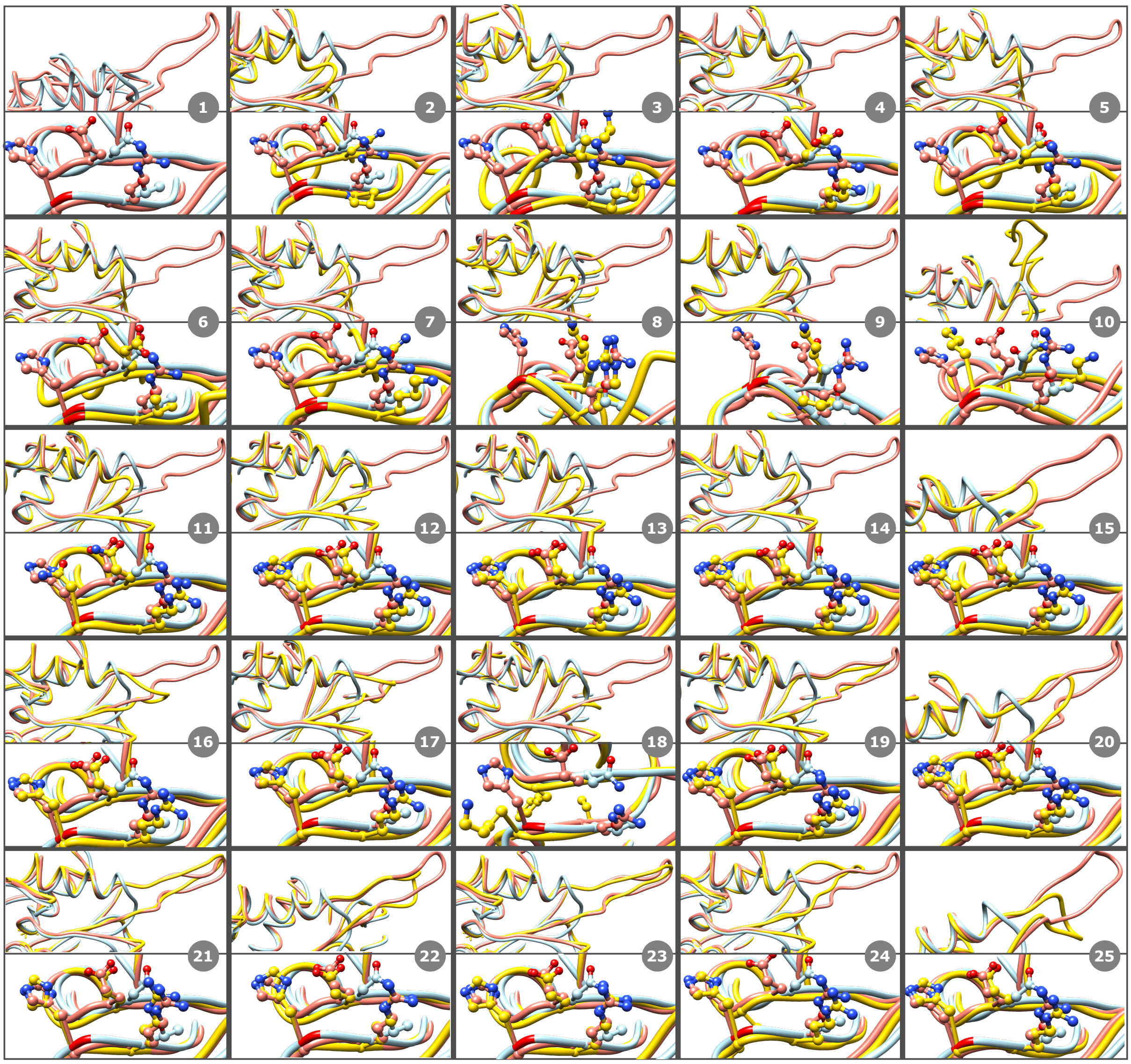

1) E.coli; 2) Methanothermobacter thermautotrophicus; 3) Methanocaldococcus jannaschii; 4) Haloarcula marismortui; 5) Korarchaeum cryptofilum (strain OPF8); 6) Cenarchaeum symbiosum (strain A); 7) Sulfolobus islandicus; 8) Odinarchaeota archaeon; 9) Candidatus Heimdallarchaeota archaeon LC_3; 10) Candidatus Heimdallarchaeota archaeon AB_125; 11) Percolomonas cosmopolites; 12) Naegleria fowleri_isoform 1; 13) Naegleria fowleri_isoform2; 14) Naegleria gruberi_isoform1; 15) Naegleria gruberi_isoform2: 16) Strigomonas oncopelti; 17) Eutreptiella gymnastica; 18) Plasmodium vivax; 19) Cafeteria roenbergensis; 20) Chenopodium quinoa; 21) Mantamonas plastica; 22) Rigifila ramosa; 23) Diphylleia rotans; 24) Dictyostelium purpureum; 25) Acanthamoeba castellani.

***Figure S11. Structural superimposition of SHMTs from all the analysed organisms*** *structural superimposition of human SHMT1 (pdb 1BJ4, pink), SHMT from E.coli (1DFO, light blue) and the one from the analysed organism (yellow). The presence or absence of the flap motif and the tetramerization residues (His 135, Arg 137, and Glu 168 in human) is highlighted. Further details are reported in table S2.*

| Analysed species | Added species | Tm (°C) |
|---|---|---|
| SHMT1 | | 58.90 ±0.04 |
| SHMT1:RNA | | 62.38±0.08 |
| SHMT1:CHOTHF5Glu | | 63.67±0.04 |
| SHMT1:CHOTHF5Glu | +RNA (0,5 μM) | 61.76±0.04 |
| SHMT1 | +ser (10 mM) | 65.99±0.05 |
| SHMT1:RNA | +ser (10 mM) | 72.28±0.27 |
| SHMT1:CHOTHF5Glu | +ser (10 mM) | 79.87±0.47 |
| SHMT1:CHOTH5Glu:ser | +RNA (0,5 μM) | 71.84±0.09 |
| SHMT1:CHOTH5Glu:ser (second peak) | +RNA (0,5 μM) | 81.80±0.99 |
| SHMT1 | +gly (10 mM) | 61.65±0.06 |
| SHMT1:RNA | +gly (10 mM) | 65.59±0.16 |
| SHMT1:CHOTHF5Glu | +gly (10 mM) | 79.12±0.10 |
| SHMT1:CHOTH5Glu:gly | +RNA (10 mM) | 78.73±0.07 |

***Table S1. Thermal melting temperatures obtained by DSF experiments****. The starting species are indicated in the left column, while the added species in the central one. The indicated temperatures are referred to the final solutions. The experimental conditions before adding the species indicated in the central column are: SHMT1 2 μM and (if present) $UTR2_{1\text{-}50}$ 0.5 μM, CHO-THF-5Glu 50 μM, serine or glycine 10 mM. The shown data were obtained by at least two individual experiments.*

| Specie | PDB | AlphaFold2 model | | Tetramer | Flap Motif | Histidine (ref. Human H135) | Arginine (ref. Human R137) | Glutamate (ref. Human E168) | RMSD Value to Human | RMSD Value to E.coli |
|---|---|---|---|---|---|---|---|---|---|---|
| E.coli | 1dfo | | | N | N | G113 | T115 | N141 | 0.902 | |
| Methanothermobacter thermautotrophicus | | AF-O27433-F1 | | N | N | G109 | P111 | Q137 | 1.05 | 0.959 |
| Methanocaldococcus jannaschii | 4bhd | | | N | N | G108 | K110 | K136 | 1.107 | 1.034 |
| Haloarcula marismortui | | AF-Q5V3D7-F1 | | N | N | G111 | K113 | E139 | 0.84 | 0.797 |
| Korarchaeum cryptofilum (strain OPF8) | | AF-B1L5K9-F1 | | N | N | G108 | V110 | D146 | 1.056 | 0.959 |
| Cenarchaeum symbiosum (strain A) | | AF-A0RYP2-F1 | | N | N | G112 | V114 | E144 | 1.006 | 0.925 |
| Sulfolobus islandicus | | AF-C3N6F2-F1 | | N | N | G109 | K111 | Q137 | 1.077 | 0.971 |
| Odinarchaeota archaeon | | AF-A0A1Q9N7E4-F1 | | N | N | G108 | R110 | K140 | 1.035 | 0.923 |
| Candidatus Heimdallarchaeota archaeon LC_3 | | AF-A0A1Q9NRJ3-F1 | | N | N | G112 | P114 | K140 | 0.915 | 0.659 |
| Candidatus Heimdallarchaeota archaeon AB_125 | | AF-A0A1Q9PFA6-F1 | | N | Y* | K112 | K114 | T140 | 0.847 | 0.691 |
| Percolomonas cosmopolitus | | AF-A0A7S1KPV0-F1 | | N | N | N126 | R128 | Q159 | 0.672 | 0.832 |
| Naegleria fowleri_1 | | AF-A0A6A5BDK2-F1 | | Y | N | H162 | R164 | E195 | 0.685 | 0.862 |
| Naegleria fowleri_2 | | AF-A0A6A5BUE9-F1 | | Y | N | H127 | R129 | E160 | 0.649 | 0.86 |
| Naegleria gruberi_1 | | AF-D2VIC8-F1 | | Y | N | H124 | R126 | E157 | 0.66 | 0.833 |
| Naegleria gruberi_2 | | AF-D2UYA5-F1 | | Y | Y | H158 | H160 | E191 | 0.676 | 0.853 |
| Strigomonas oncopelti | | AF-U5KMX5-F1 | | Y | Y* | H116 | R118 | E149 | 0.754 | 0.843 |
| Eutreptiella gymnastica | | AF-A0A7S1NGR6-F1 | | Y | Y* | H125 | R127 | E159 | 0.713 | 0.854 |
| Plasmodium vivax | 4tn4 | | | N | N | K116 | I119 | E148 | 0.74 | 0.934 |
| Cafeteria roenbergensis | | AF-A0A5A8C385-F1 | | Y | Y | H146 | H148 | E179 | 0.671 | 0.896 |
| Chenopodium quinoa | | AF-A0A803LBD8-F1 | | Y | Y* | H197 | R199 | E230 | 0.622 | 0.894 |
| Mantamonas plastica | | | generated models | Y | Y | H157 | H159 | E190 | 0,446 | 0.921 |
| Rigifila ramosa | | | generated models | Y | Y | H127 | R129 | E160 | 0.411 | 0.909 |
| Diphylleia rotans | | | generated models | Y | Y | H142 | R144 | E175 | 0.378 | 0.892 |
| Dictyostelium purpureum | | AF-F0ZDZ0-F1 | | Y | Y | H144 | R146 | E176 | 0.653 | 0.862 |
| Acanthamoeba castellani | | AF-L8HDT1-F1 | | Y | Y | H120 | R122 | E153 | 0.639 | 0.833 |

***Table S2. Phylogenetic analysis.*** *All the analysed species and their PDB or AlphaFold 2 models codes are indicated in this table. The presence of the tetramerization residues (H135, R137, E168 in humans) is reported in columns 7-9. The RMSD values of Cα for each species are indicated in the last two columns. Flap motifs that are shifted and/or shorter than the human are indicated as Y*.*
